# Surface display of designer protein scaffolds on genome-reduced strains of *Pseudomonas putida*

**DOI:** 10.1101/2020.05.13.093500

**Authors:** Pavel Dvořák, Edward A. Bayer, Víctor de Lorenzo

## Abstract

The bacterium *Pseudomonas putida* KT2440 is gaining considerable interest as a microbial platform for biotechnological valorization of polymeric organic materials, such as waste lignocellulose or plastics. However, *P. putida* on its own cannot make much use of such complex substrates, mainly because it lacks an efficient extracellular depolymerizing apparatus. We seek to meet this challenge by adopting a recombinant cellulosome strategy for this attractive host. Here, we report an essential step in this endeavor – a display of designer enzyme-anchoring protein “scaffoldins”, encompassing cohesin binding domains from divergent cellulolytic bacterial species on the *P. putida* surface. Two *P. putida* chassis strains, EM42 and EM371, with streamlined genomes and substantial differences in the composition of the outer membrane were employed in this study. Scaffoldin variants were delivered to their surface with one of four tested autotransporter systems (Ag43 from *Escherichia coli*), and the efficient display was confirmed by extracellular attachment of chimeric β-glucosidase and fluorescent proteins. Our results highlight the importance of cell surface engineering for display of recombinant proteins in Gram-negative bacteria and pave the way towards designer cellulosome strategies, tailored for *P. putida*.

## Introduction

Polymeric organic materials such as waste lignocellulose or plastics represent a potentially inexhaustible source of cheap carbon and energy for biotechnology and synthetic biology enterprises.^1,2^ Adoption of these recalcitrant feedstocks for bioproduction of valuable chemicals nonetheless requires the employment of microbial hosts with a suite of properties that would allow them to perform complex biocatalytic conversions efficiently even under harsh conditions of industrial processes. Such microorganisms are currently not available but can be obtained by engineering suitable robust platform strains.

*Pseudomonas putida* KT2440, a popular Gram-negative bacterial workhorse, definitely fulfills some of the crucial criteria to become a host of choice for biotechnological upcycling of polymeric wastes. It has been recently engineered for utilization and valorization of several plant biomass-derived sugars,^3–5^ lignin-born aromatic chemicals^6,7^ or even products of synthetic plastic degradation.^8^ It was also demonstrated that this bacterium can process oligomeric carbohydrates^4,9^ as well as co-utilize hexose and pentose sugars – glucose and xylose - and consume simultaneously glucose and an aromatic substrate with a lack of diauxia.^4,10^ These and other characteristics including safety status,^11^ rapid growth and low nutritional demand,^12^ considerable resistance to inhibitory chemicals,^13,14^ its employment in large-scale fermentations for production of value-added chemicals,^15^ or its compliance to genetic manipulations and the available palette of engineering tools,^16–18^ make *P. putida* an attractive candidate for the demanding biotechnological task sketched above. However, *P. putida*, same as the majority of other domesticated microbial platforms, lacks efficient extracellular depolymerizing apparatus and cannot degrade complex recalcitrant substrates alone.

The most efficient natural polymer degraders known to date, cellulolytic bacteria such as *Clostridium thermocellum*, display on their surface cellulosomes – remarkable nanomachines composed of scaffoldin proteins that attach and orchestrate multiple carbohydrate-active enzymes on the cell surface.^19^ The binding of cellulases to scaffoldins is mediated by strong (K_D_ ∼ 10^−9^ – 10^−10^ M) highly specific non-covalent interactions between cohesin and dockerin binding domains.^20^ Extensive H-bond networks provide cohesin-dockerin pairs the firmness which reaches half of the mechanical rupture strength of a covalent bond and surpasses antigen-antibody binding.^21,22^ Concerted action of dockerin-tagged cellulases clustered on the cell surface through interactions with cohesins in scaffoldin proteins can enhance cellulose degradation up to 50-fold when compared with free enzymes.^23^

Natural cellulosome producers (*e.g., C. thermocellum, Bacteroides cellulosolvens, Acetivibrio cellulolyticus*) are often difficult to genetically manipulate or cultivate, and their cellulosomes are large and complicated.^24^ Hence, smaller synthetic *designer cellulosomes* or *minicellulosomes* have been assembled either *in vitro* using purified components^25–27^ or *in vivo* on the surface of a suitable microbial host.^28–32^ A fundamental prerequisite for successful minicellulosome assembly is a display of scaffoldin proteins on the surface of a target host. Truncated forms of native scaffoldins or designer hybrid scaffoldins with cohesin domains from diverse cellulolytic organisms were delivered to the cell surface of a recombinant *Saccharomyces cerevisiae*,^28,33^ *Bacillus subtilis*,^31^ *Clostridium acetobutylicum*,^34^ *Lactobacillus plantarum*^32^ or *Lactococcus lactis*.^35^ However, efficient expression and secretion of cellulosome components in a phylogenetically distant Gram-negative host with different codon usage, G+C content in a genome, a complicated two-layer structure of cell wall, and crowded cell surface remains to be very challenging. Not surprisingly, surface display of scaffoldins and subsequent *in vivo* assembly of a designer cellulosome of any size has not yet been reported in a biotechnologically relevant Gram-negative bacterium. Thus far, display or secretion of individual depolymerizing enzymes has been achieved in engineered *E. coli*, allowing it to grow on cellooligosaccharides or produce a limited quantity of biofuels from plant biomass.^36–39^ In the case of *P. putida*, single cellulases from *Ruminiclostridium thermocellum* and hemicellulases from *Bacillus subtilis* were displayed on the surface of recombinant strains mixed to form designer co-cultures of resting cells.^40,41^ The joint activities of these enzymes in cell suspensions of high cell densities resulted in the production of small quantities of glucose^40^ or xylose^41^ from filter paper or arabinoxylan, respectively, which were nonetheless insufficient to support the growth of the host bacterium. Adoption of the designer cellulosome approach can result in more efficient saccharification of (hemi)cellulose when compared to the strategy which utilizes cellulases displayed separately on several strains.^42^ However, to test that hypothesis also in a selected Gram-negative bacterial host, an efficient display of scaffoldins for docking of cellulases on its surface must first be achieved.

In this study, we aimed to display structurally distinct variants of a designer miniscaffoldin on the surface of recently introduced genome-reduced *P. putida* strains designated EM42 and EM371 (**Fig 1 and Fig. S1**). The two strains were adopted mainly due to the substantial differences in presence of diverse outer membrane structures and complexity of the bacterial surface which can impose structural constraints for displayed proteins.^43^ Strain EM42 possesses eleven non-adjacent genomic deletions (300 genes, ∼ 4.3 % of the whole genome) that were shown to improve expression of heterologous genes and enhanced biotechnological potential of this *P. putida* KT2440 derivative.^4,44,45^ Except for missing flagellum, the cell surface of *P. putida* EM42 resembles that of the wild-type strain KT2440. In the strain EM371, on contrary, most of the non-essential outer membrane structures that are used by bacteria to coordinate motion (flagella) or to develop biofilms and interact with their surroundings (*e.g*., fimbriae, pili, curli, adhesins, exopolysaccharides, lipopolysaccharides) were eliminated (230 genes, ∼ 4.7 % of the entire genome).^46^ The “shaved” surface endowed *P. putida* EM371 with properties potentially beneficial for extracellular recombinant protein production and facilitated downstream processing.^46^ Here, expression and display of scaffoldin variants on the surface of EM42 and EM371, *via* one of the four tested autotransporter systems, allowed: (i) addressing important questions on how does the character of secreted molecules and cellular exterior contribute to cohesin-dockerin interactions and (ii) comparison of the capability of the two biotechnologically relevant *P. putida* strains to become Gram-negative bacterial platforms for *in vivo* assembly of surface-exposed designer depolymerizing nanomachines.

**Figure 1.**
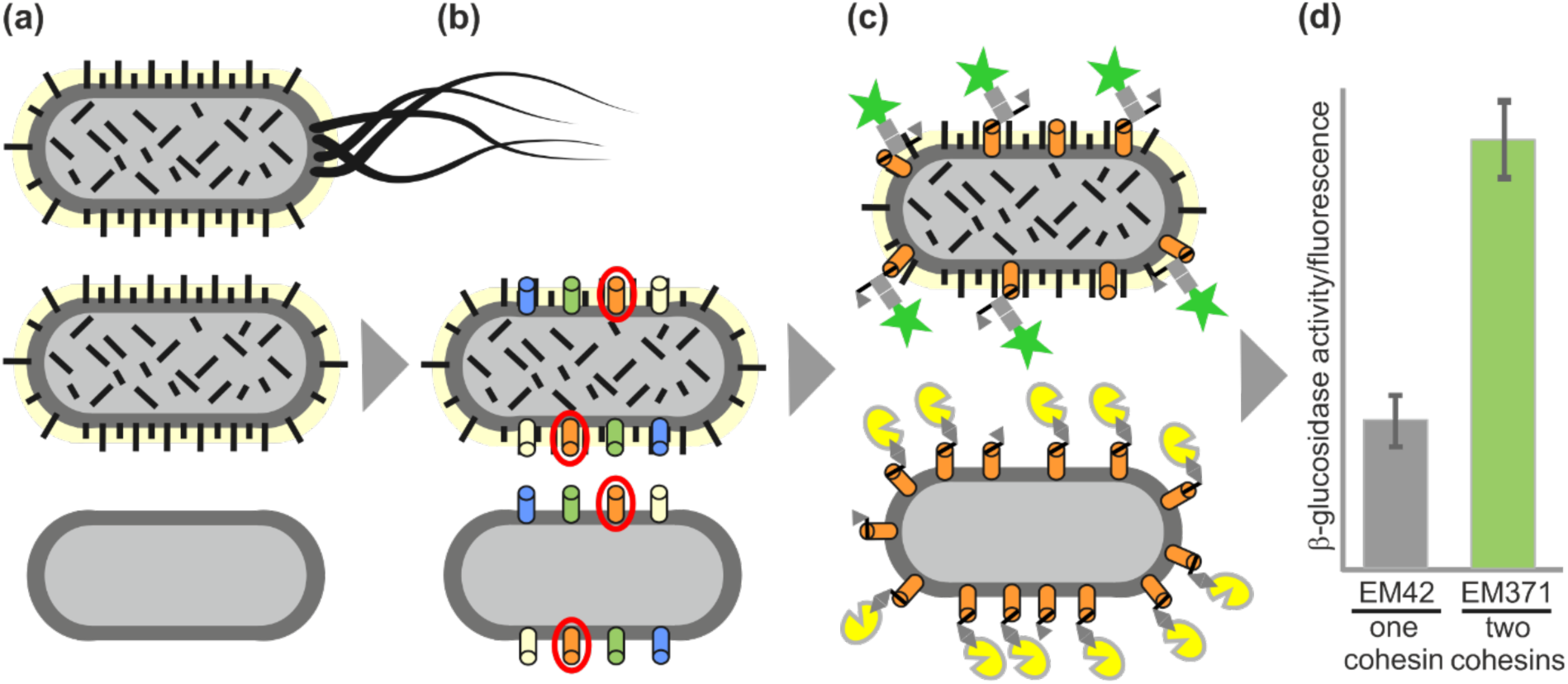
Schematic overview of the actions: Engineering *Pseudomonas putida* for surface display of cohesin-containing designer protein scaffoldins. **(a)** Two derivatives of *P. putida* KT2440 (top graphics), strains EM42 and EM371, with streamlined genomes lacking de-stabilizing genetic elements and certain surface structures, respectively, were employed in this study. **(b)** Four type V secretion systems were tested in the target host and the best-performing autotransporter was selected for further work. **(c)**+**(d)** Single- or two-cohesin scaffoldins were displayed on EM42 and EM371 surface and the efficiency of the binding of dockerin-tagged recombinant proteins (β-glucosidase or fluorescent proteins) to the cellular surface was evaluated and quantified.

## Results

### Preparation of cohesin-containing scaffoldin variants and dockerin-tagged chimeric proteins

Scaffoldin Scaf19L assembled by Vazana and co-workers^47^ was initially adopted for this study (**Fig. 2a, Supplementary sequences in SI**). This chimaeric construct contains the N-terminal carbohydrate-binding module (CBM) from *Clostridium thermocellum* and three cohesin domains, designated here AcCoh, CtCoh, and BcCoh, from *Acetivibrio cellulolyticus, C. thermocellum*, and *Bacteroides cellulosolvens*, respectively (for details see Material and methods section). The cohesins and CBM are interconnected with 27 – 35 amino acid (aa) long flexible linkers, which were shown to have a positive effect on the overall activity of a synthetic minicellulosome assembled *in vitro*.^47^ The scaffoldin gene was subcloned into the pSEVA238 plasmid with inducible XylS/Pm expression system^48^, and the construct was inserted into *Pseudomonas putida* EM42. However, induction of the gene expression resulted in reduced host’s fitness (**Fig. S2a**). Western blot analysis of the soluble and insoluble fractions of the cell lysate revealed limited Scaf19L solubility and its susceptibility to proteolytic cleavage (**Fig. S2b**). We argued that these problems could be caused by a discrepancy between the mean GC content (38 %) and codon distribution in the chimeric gene and in the genome of the host organism (mean GC content in *P. putida* genome is 62 %^49^). The codon adaptation index (CAI), calculated for *scaf19L* gene and *P. putida* KT2440 host using the on-line tool JCat^50^, was very low (0.10; where CAI of 1.0 signifies the best match), which indicated that gene toxicity could indeed have reflected codon bias. Hence, the *scaf19L* gene was synthesized and codon optimized for expression in *P. putida*. Heterologous expression of the resultant optimized gene *scaf19LKT* with increased GC content (57 %) had no negative effect on *P. putida* EM42 growth (**Fig. S2a**). Most of the Scaf19LKT protein was produced in a soluble form (>75 % compared to < 50 % before optimization), and proteolysis was not observed (**Fig. S2b**). These results show that GC content and codon usage are two key sequence features that must be taken into consideration for design of synthetic cellulosome components from phylogenetically distant species to be expressed in *P. putida*.

**Figure 2.**
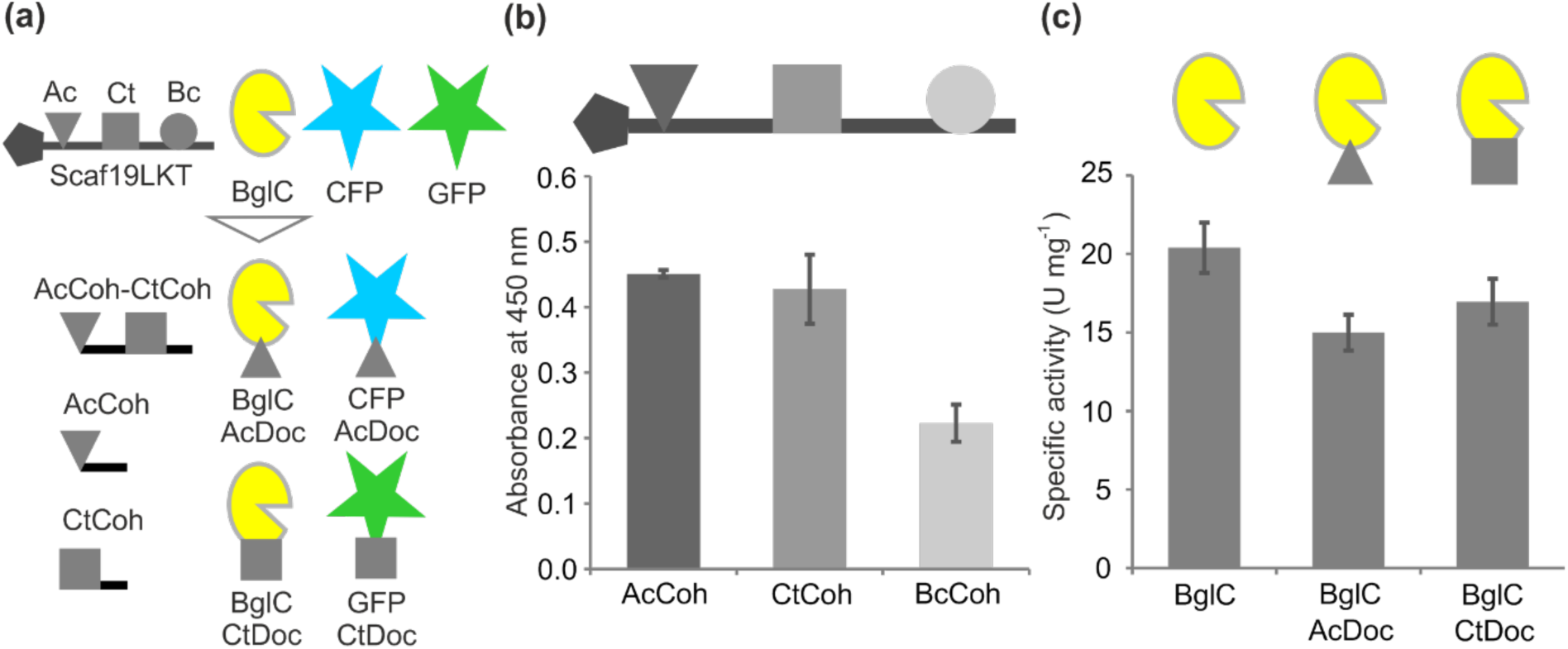
Recombinant proteins used in this study and initial evaluation of their function. **(a)** Three truncated variants of synthetic scaffoldin gene *scaf19LKT* (for detailed description see Materials and Methods or work of Vazana and co-authors^47^), codon-optimized for expression in *Pseudomonas putida*, were prepared and adopted for this study: two variants with either single CtCoh (square symbol) or AcCoh (triangle symbol) cohesin from *Clostridium thermocellum* or *Acetivibrio cellulolyticus*, respectively, and a two-cohesin scaffoldin with both AcCoh and CtCoh interconnected with a 29 amino acid-long linker. Furthermore, two chimeric variants of β-glucosidase BglC from *Thermobifida fusca* with CtDoc or AcDoc dockerin matching complimentary cohesins were constructed, as well as a recombinant monomeric superfolder green fluorescent protein (designated here simply as GFP) with CtDoc and mCerulean fluorescent protein (designated here as CFP) with AcDoc. **(b)** ELISA-based (enzyme-linked immunosorbent assay) verification of proper folding and function of AcCoh, CtCoh, and BcCoh in the Scaf19LKT scaffoldin produced in *P. putida* EM42. Data are shown as mean ± SD from two biological replicates, each conducted with two technical replicates. **(c)** Comparison of activities of purified dockerin-tagged BglC variants produced in *P. putida* EM42 with activity of wild-type BglC. Data are shown as mean ± SD from three biological replicates.

The functionality of cohesins in Scaf19LKT produced in *P. putida* EM42 was then analyzed using an enzyme-linked immunosorbent assay (ELISA)-based binding assay.^47^ The assay was based on binding of recombinant variants of *Geobacillus sp.* endo-1,4-β-xylanase fused to dockerins from *A. cellulolyticus, C. thermocellum*, or *B. cellulosolvens* (named here AcDoc, CtDoc, or BcDoc, respectively) to respective cohesins in Scaf19LKT molecules in *P. putida* EM42 cell-free extract and subsequent detection of the assembled complexes with anti-xylanase antibody. **Figure 2b** shows that all three cohesins in Scaf19LKT produced in *P. putida* were able to bind their respective dockerins but not with equal efficiency. AcCoh-AcDoc and CtCoh-CtDoc pairs provided twice as much signal as the BcCoh-BcDoc combination, which signified a possible tighter binding of the former two pairs. We therefore selected these pairs for experiments in the current study. Three truncated variants of *scaf19LKT* gene encoding scaffoldins AcCoh, CtCoh, and AcCoh-CtCoh with a single cohesin or with two cohesins were prepared for surface display on *P. putida* EM42 and EM371 strains (**Fig. 2a**).

We then aimed at the assembly of dockerin-tagged reporter proteins that could be employed in a rapid robust assay for the detection of displayed scaffoldins on the surface of the target host cells. Displayed cohesins can be detected and quantified by ELISA-based protocols^32,47^ or by assays with dockerin-tagged enzymes, such as β-glucuronidase.^35^ The latter approach was adopted in this study. β-Glucosidase (EC 3.2. 1.21) BglC from *Thermobifida fusca* was previously shown to be functionally expressed in *P. putida* EM42 up to 30 % of the total soluble protein.^4^ Measurement of its hydrolytic activity with synthetic *p*-nitrophenyl-β-D-glucopyranoside (pNPG) substrate is simple and fast. We modified this enzyme by fusing it on its C terminus with AcDoc or CtDoc dockerin (**Fig. 2a**). Two chimeras and wild-type BglC with a polyhistidine tag were produced in *P. putida* EM42, purified by immobilized metal affinity chromatography (**Fig. S3a – S3c**), and specific activities of the three enzymes with pNPG were determined. As shown in **Figure 2b**, enzyme fusion to neither of the two dockerins reduced the activity substantially. Activities of BglC-AcDoc and BglC-CtDoc reached 74 % and 83 % of the wild-type activity, respectively. The lower performance of BglC-AcDoc compared to BglC-CtDoc can be attributed to the lower expression and consequent also lower purity of this chimera (70 % and 85 % protein purity of BglC-AcDoc and BglC-CtDoc, respectively, was estimated from sodium dodecyl sulfate (SDS) polyacrylamide gels using ImageJ Gel Densitometry Tool, **Fig. S3b and S3c**). Dockerin-tagged variants of cyan fluorescent protein mCerulean and monomeric superfolder green fluorescent protein, abbreviated here as CFP-AcDoc and GFP-CtDoc, respectively, were prepared according to BglC chimeras for spectroscopic and microscopic confirmation of displayed scaffoldins (**Fig. 2a**). Chimeric fluorophores were produced in *Escherichia coli* BL21-Gold (DE3) and purified by affinity chromatography (**Fig. S3D and S3E**).

### Selection of an optimal autotransporter system for display of scaffoldins on the surface of P. putida EM42 and EM371 strains

Monomeric type V secretion pathway proteins known as autotransporters have been extensively used for decorating surfaces of Gram-negative bacteria with recombinant proteins.^51,52^ They became popular mainly due to their simplicity (a single gene encodes all three domains needed for display – a signal peptide, a surface-exposed passenger, and a transmembrane β-domain), high display efficiencies reaching in certain cases 10^4^ - 10^5^ enzyme molecules per cell, and relatively low toxicity of recombinant variants towards a bacterial host.^53,54^ Three autotransporters, namely EhaA from enterohemorrhagic *E. coli*,^55^ EstP from *P. putida*,^56^ and immunoglobulin A (IgA) protease from *Neisseria gonorroheae*^57^ were used with some success for a passenger export in the KT2440 strain or its derivative, but, in general, the reports on recombinant protein surface display in this host are scarce.

It is often highlighted in the scientific literature that the accurate prediction of a secretion system, functioning with a given passenger protein in a selected bacterial host, is unlikely.^58^ Hence, it is desirable to test at least several candidate systems. Here, we sought to evaluate four different autotransportes for display of designer scaffoldins in *P. putida*. Three previously constructed systems, including translocator domains (β-barrels and α-helix linkers) of the IgA protease from *N. gonorroheae*,^59^ antigen 43 (Ag43) from *E. coli*,^38,60^ and EaeA intimin (Int, inverse autotransporter) from *E. coli*^61,62^ were adopted for this study (**Fig. 3a, Supplementary Table S1, Material and methods**). Moreover, *P. putida* EstP esterase (PP_0418) translocator sequence, encoding the β-barrel domain with a spanning α-helical linker complemented by a synthetic multi-cloning site and a native signal peptide sequence, was synthesized and added to the list for testing. IgA and intimin were already available in our laboratory, cloned in pSEVA238 plasmid. Ag43 and EstP autotransporter genes were subcloned into this vector from a provided^38^ and a delivery plasmid, respectively. The EstP autotransporter was nonetheless soon excluded from the list, because its expression in *P. putida* EM42 appeared to be extremely toxic for the host (**Fig. S4**). Such a strong toxic effect might be attributed to the burden caused by overproduction of the native protein with efficient secretion signal and subsequent distortion of the cell membrane due to the high number of integrating β-barrels.^63^ Indeed, we observed formation of flocks in the culture after induction, which is a common stress response mechanism, observed, *e.g.*, in cells with overexpressed porins or other membrane proteins.^5,64^ Overproduced autotransporter molecules can also exhaust secretion machinery – namely Sec and BAM systems - required for the export of proteins necessary for cell growth and maintenance.^51,65^ Induction of expression of the remaining three autotransporters had no or negligible effect on the host’s viability (**Fig. S4**). All these systems were therefore selected for further testing.

**Figure 3.**
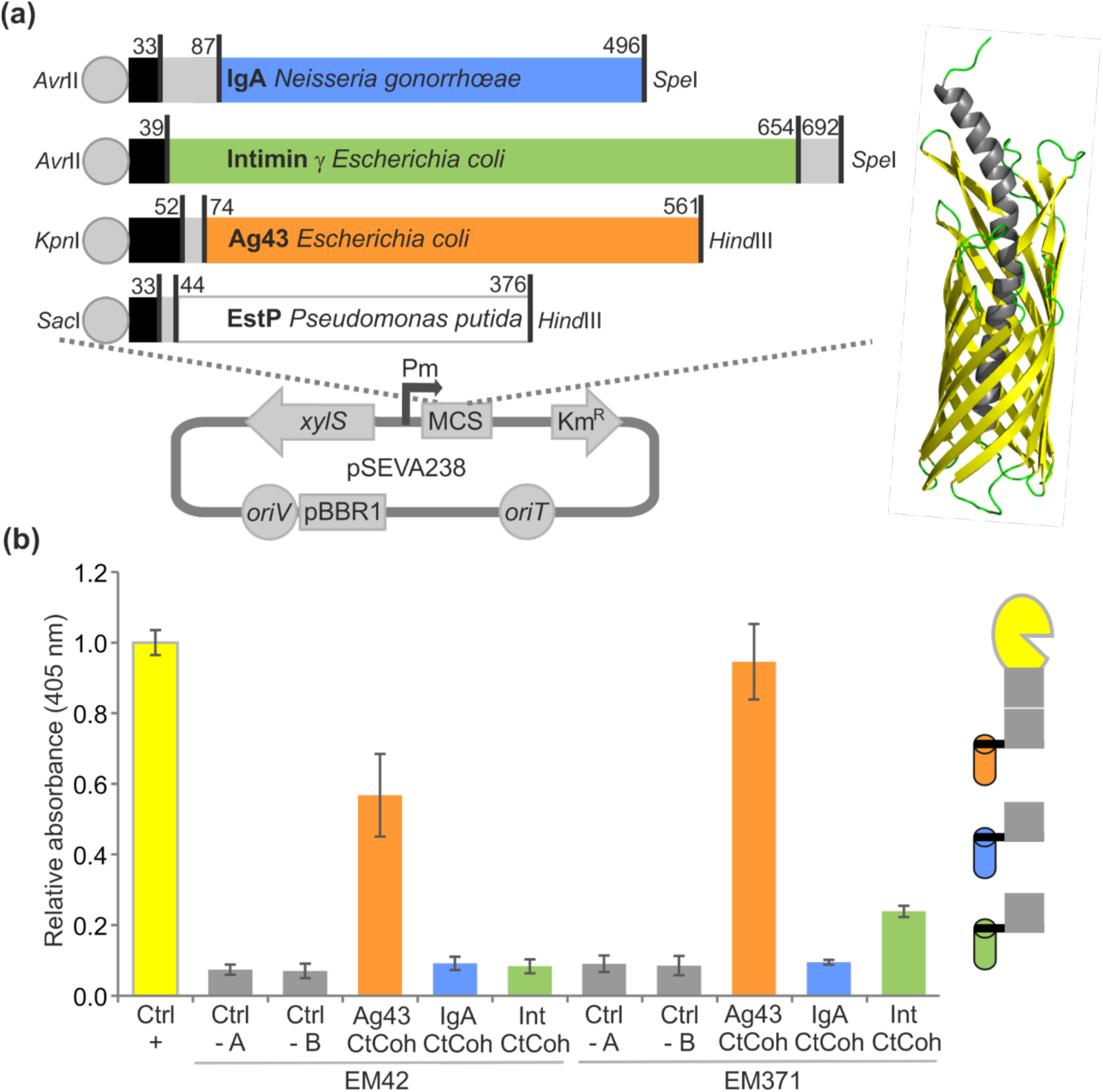
Selection of the type V secretion system for display of scaffoldins on the surface of *Pseudomonas putida* EM42 and EM371 strains. **(a)** Cloning of genes encoding translocation β-barrel domain and part of the α-helix passenger (model of EstP autotransporter prepared by iTasser^85^ is shown as a representative example) of four autotransporter systems from three different Gram-negative bacteria into the marked restriction sites of the pSEVA238 plasmid polylinker. All four genes were preceded by a synthetic ribosome binding site (grey sphere) and contained a signal peptide sequence (in black) and their own multiple cloning site (in grey). The length of the individual segments in the amino acid sequence of a given autotransporter is shown. **(b)** Test display of CtCoh cohesin on the surface of EM42 and EM371 strains with three selected secretion systems (Ag43, IgA, and intimin). The cells displaying CtCoh were mixed with purified BglC-CtDoc, washed and incubated with *p*-nitrophenyl-β-D-glucopyranoside substrate. End-point absorbance of the reaction product *p*-nitrophenol (corresponds to β-glucosidase activity of the whole cells with anchored BglC) was determined and compared with the absorbance measured for purified BglC-CtDoc of defined concentration in the reaction mixture (Ctrl+). EM42 and EM371 cells with empty pSEVA238 plasmid mixed with BglC-CtDoc (Ctrl-A), and EM42 and EM371 pSEVA238_*ag43AT-CtCoh* cells mixed with wild-type BglC (Ctrl-B) were used as negative controls. Data are shown as mean ± SD from at least three independent experiments, each conducted in two technical replicates.

In the next step, the codon-optimized gene encoding CtCoh was subcloned into the polylinker of *igAAT, ag43AT*, and *intAT*, and the three autotransporters were tested for display of the single-cohesin scaffoldin of the theoretical molecular weight of 16.2 kDa on the surface of EM42 and EM371 strains. Cells displaying scaffoldin were added with an excess of purified BglC-CtDoc (∼1.0×10^5^ molecules per cell, the value was determined as described in the Materials and methods section), washed, and their β-glucosidase activity was determined by measuring the end-point absorbance of the reaction product *p*-nitrophenol released after hydrolysis of the pNPG substrate (**Fig. 3b**). The absorbance of the supernatant fluids from the reactions with whole cells was related to the absorbance of the mixture with purified BglC-CtDoc, which served as a positive control. An obvious advantage of such an approach is that all measured activity can be attributed only to the chimeric enzyme molecules attached from outside of the surface of the intact cells that passed cycles of centrifugation and washing. This feature is especially valuable when considering the fact that whole-cell ELISA and proteinase accessibility assays, frequently used to evaluate the efficiency of recombinant protein display, are prone to false-positive results.^54,66^

**Figure 3b** reveals that only the cells expressing Ag43 autotransporter showed high β-glucosidase activity, which, in the case of the EM371 recombinant, reached the level of the pure enzyme control. Some activity was also detected with EM371 cells displaying scaffoldin *via* intimin, but their EM42 counterparts were not able to attach dockerin-tagged BglC to their surface. Absorbance measured in supernatant fractions from reactions with the EM42 pSEVA238_*intAT-CtCoh* recombinant did not surpass that of the negative controls (**Fig. 3b**). The same was also true for both recombinants (either EM42 or EM371) expressing *igAAT*, which suggests that this autotransporter was the least efficient in CtCoh display out of the three tested candidates. SDS polyacrylamide gel electrophoresis (SDS-PAGE) and western blot analysis of the cell lysates (**Fig. S5a and S5b**) confirmed the good expression of the *ag43AT-ctCoh* gene in both EM42 and EM371 recombinants. The chimera comprised around 3 % of the total cellular protein, as determined using GS-800 Calibrated Densitometer (Bio-Rad). In contrast, expression of *intAT-ctCoh* was not detected on SDS-PAGE gel in either of the two lysates, and only a very pale barely detectable band was identified among the blotted proteins of the EM371 recombinant (**Fig. S5a and S5b**). Hence, the malfunctioning of this autotransporter system in the current study can be attributed to the poor expression of the construct. The same conclusion cannot be made for IgAAT-CtCoh because its production was detectable in EM42 and EM371 lysates, both on the SDS-PAGE gel and on the blotting membrane. However, no signal was seen with whole pre-induced EM42 and EM371 cells bearing the *igAAT-ctCoh* gene after their incubation with HRP-conjugated anti-6xHis tag antibody during the dot blot analysis (**Fig. S5c**). Hence, one possible explanation of the IgAAT-CtCoh malfunctioning is that the chimera was expressed, but it was not functionally displayed on the *P. putida* surface, perhaps due to improper folding during its transport and maturation. It is worth noting that *P. putida*, empowered with the IgA translocator construct identical to the one used here, was able to secrete detectable quantities of metallothioneins or eukaryotic leucine zippers in the former works of Valls et al.^67^ and Martínez-García and co-workers, respectively. Such discrepancy supports the repeatedly certified observation that the ability of a certain autotransporter to export a passenger of choice in a given host bacterium cannot be predicted with confidence prior to experimental verification.^54,58^

Dot blot analysis of whole pre-induced cells also confirmed the display of CtCoh *via* the Ag43 autotransporter in the EM371 strain (**Fig. S5c**). None of the EM42 strains, including EM42 with the Ag43AT-CtCoh construct, showed a luminescence signal. We hypothesize that this could be ascribed to the location of the 6xHis tag between the CtCoh and Ag43AT molecules and its poor accessibility for antibody on the crowded surface of the EM42 cells. Taken together, the results discussed above identified Ag43 as a promising secretion system for surface display of cohesin binding domains in *P. putida*.

Native Ag43 (Uniprot ID: P39180) possesses an N-terminal signal peptide (aa 0-52), a surface-exposed N-proximal β-helical α passenger domain (aa 53-551) which is responsible for an autoaggregation phenotype in *E. coli*, and a C-terminal β-barrel domain (aa 552-1039) whose substantial part, except for an autochaperone domain that facilitates folding of the passenger domain, is buried in the outer membrane.^68^ Under normal circumstances, the α domain is cleaved during the secretion process by an internal protease motif and remains attached to the cell surface by noncovalent interactions that can be disrupted, *e.g*., by heat treatment or osmotic shock.^69^ The Ag43 autotransporter employed in this study lacks the whole α domain, including the aspartyl protease active site which was removed and substituted with a polylinker for cloning of a passenger of choice.^38^ The passenger protein thus remains covalently attached to the cell surface.

### Binding of dockerin-tagged β-glucosidase and fluorescent proteins on P. putida EM42 and EM371 cells displaying scaffoldin variants

Cellulase BglC-CtDoc was successfully attached to *P. putida* cells expressing Ag43 autotransporter with a single-cohesin scaffoldin. But how is such directed interaction on bacterial surface influenced by the character and size of the displayed scaffoldin? And how does the character of a host cell’s exterior contribute to such bonding? We aimed to answer these questions in the following part of our study. Genes encoding two remaining scaffoldin variants – single-cohesin AcCoh (theoretical Mw=16.5 kDa) and two-cohesin AcCoh-CtCoh (theoretical Mw=34.6 kDa) – were subcloned separately into the polylinker of *ag43AT* in the pSEVA238 vector, and the resulting constructs were transferred to “hairy” EM42 and “naked” EM371 cells with preserved and eliminated outer membrane structures, respectively. Surface display of the structurally distinct AcCoh, CtCoh, and AcCoh-CtCoh scaffoldins on the *P. putida* recombinants was evaluated by attachment of dockerin-tagged β-glucosidase variants BglC-AcDoc and BglC-CtDoc and chimeric fluorophores CFP-AcDoc and GFP-CtDoc (**Fig. 4**). All purified recombinant proteins were added to the cells in high excess (∼1.0×10^5^ β-glucosidase and ∼1.8×10^5^ fluorophore molecules per cell) to secure saturation of cohesin domains on the surface with the respective dockerins.

**Figure 4.**
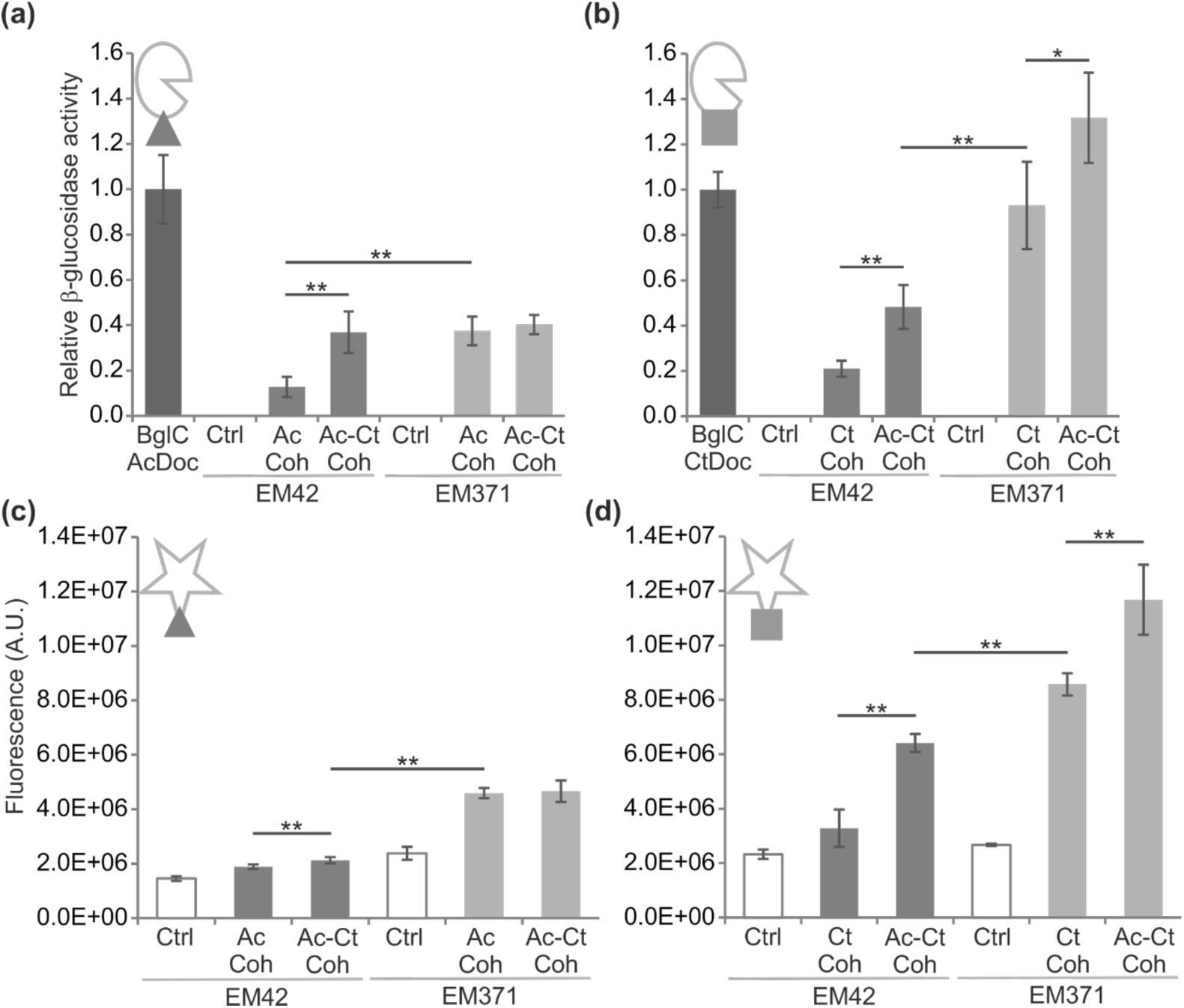
Binding of dockerin-tagged β-glucosidase and fluorescent proteins on *Pseudomonas putida* EM42 and EM371 cells displaying designer scaffoldins. **(a)** + **(b)** Binding of BglC-AcDoc and BglC-CtDoc, respectively, to EM42 and EM371 cells displaying AcCoh, CtCoh, or AcCoh-CtCoh scaffoldins. β-glucosidase activities were measured with whole cells and related to the activity of purified BglC-AcDoc (4.64 ± 0.71 U) or BglC-CtDoc (4.91 ± 0.39 U). EM42 and EM371 cells with the pSEVA238b_*ag43AT* plasmid were used as controls (Ctrl). Data are shown as mean ± SD from at least two independent experiments, each conducted in two to three biological replicates. **(c)** + **(d)** Binding of CFP-AcDoc and GFP-CtDoc, respectively, to EM42 and EM371 cells displaying AcCoh, CtCoh, or AcCoh-CtCoh scaffoldins. CFP and GFP fluorescence was measured at 440/500 nm and 475/515 nm, respectively. EM42 and EM371 cells with pSEVA238b_*ag43AT* plasmid were used as controls (Ctrl). Data are shown as mean ± SD from three biological replicates, each conducted in two technical replicates. Asterisks denote significance in difference in between two means at P < 0.05 (*) or P < 0.01 (**).

Whole-cell activities of β-glucosidase were quantitatively determined in defined time intervals and related to the activity of the corresponding purified recombinant enzyme (**Fig. 4a and 4b, Material and methods**). Fluorescence of *P. putida* cells with surface-attached CFP or GFP was measured in microtiter plate format (**Fig. 4c and 4d**). The results confirmed that both cohesin domains were accessible on the cell surface and functional in binding their respective dockerins. Similar trends in the binding efficiency of dockerin-tagged proteins to the cells with displayed cohesins were observed in both assay types. EM371 cells bound more dockerin-tagged proteins and showed significantly higher β-glucosidase activity and fluorescence (P < 0.01) than EM42 recombinants in almost all tested scenarios. This could not be ascribed to the better expression of autotransporter-scaffoldin chimeras in the naked strain. On the contrary, the expression levels of all three constructs were approximately 25 % lower in EM371 than in EM42, as judged by densitometric analysis of the SDS-PAGE gel with samples of cell lysates (**Figs. S6**). Hence, we assume that better display or accessibility of the exported scaffoldins on the surface of the naked strain is the correct explanation. Deletion mutants of *E. coli* BL21(DE3) lacking several abundant but non-essential outer membrane proteins were previously shown to be excellent hosts for overexpression of heterologous membrane β-barrel proteins, presumably because targeted knockouts relieved some of the burden on the Sec and BAM secretion machineries.^51^ Moreover, removed surface structures leave more space for heterologous transporters and their passengers, which can make them better accessible for their binding partners.^64^

Out of the two tested cohesin-dockerin pairs, AcCoh and AcDoc showed lower binding affinity in both EM42 and EM371 strains (**Fig. 4**). This observation does not match the former outcome of the ELISA assay which indicated equally strong bonds in the two pairs (**Fig. 2b**). However, the current experiments were performed with whole cells, not cell-free extracts as in the case of ELISA, and it is possible that AcCoh architecture was affected during the secretion process in the non-native host. In the case of fluorescence measurements (**Fig. 4c and 4d**), the dimmer signals from the cells decorated with AcCoh + CFP-AcDoc complexes could partially stem also from lower brightness of the cyan fluorophore when compared with GFP.^70^ However, the most probable cause of the observed phenomenon is that the cohesin-dockerin pair from the mesophilic *Acetivibrio cellulolyticus* is less stable and more prone to disruption during the experimental treatment (including several cycles of cell centrifugation and washing) than the binding domains from thermophilic *Clostridium thermocellum*. As shown recently by Gunnoo and co-workers^22^, who performed molecular dynamics simulations with the very same cohesin-dockerin pairs, the *C. thermocellum* system presents a stronger hydrogen bond network and higher binding affinities than the *A. cellulolyticus* complex even at ambient temperature. These theoretical calculations together with our experimental data indicate that the binding domains from thermophilic cellulolytic bacteria might be a better choice for assembly of stable scaffoldin-enzyme interactions also on the surface of mesophilic microbial hosts.

Another conclusion which deserves attention is that *P. putida* cells displaying larger AcCoh-CtCoh scaffoldins were able to bind more enzyme or fluorophore molecules than recombinants decorated with single cohesins (**Fig. 4**). This trend was especially pronounced with *P. putida* EM42 pSEVA238_*AcCoh-CtCoh*. The strain showed 2.9-fold or 2.3-fold higher β-glucosidase activity (**Fig. 4a and 4b**) and 1.5-fold or 4.3-fold enhanced fluorescence (**Fig. 4c and 4d**) when compared with its AcCoh- or CtCoh-exporting counterparts, respectively. This somewhat counterintuitive observation is in agreement with the study of Wieczorek and Martin^35^ who described the same phenomenon for the secretion of designer scaffoldins in the Gram-positive bacterium *Lactococcus lactis*. Larger molecules may make more space for themselves on the “bushy” surface of EM42 strain and are thus better accessible for binding partners than buried single cohesins. This hypothesis is supported by the fact that such a vast difference between the display of single- and two-cohesin scaffoldin was not observed with shaved EM371 recombinants (**Fig. 4**). In these, both smaller and larger scaffoldins were equally (**Figs. 4a and 4c**) or similarly (**Figs 4b and 4d**, the difference is 1.4- and 1.5-fold) accessible for AcDoc- and CtDoc-tagged proteins, respectively. The higher accessibility of the two-cohesin scaffoldin in the EM42 recombinant was certainly not the outcome of its better expression. On the contrary, AcCoh-CtCoh was the least expressed construct from all three tested scaffoldin variants both in EM42 and EM371 (**Fig. S6**).

The aforementioned phenomena were re-confirmed by observing the cells with surface-docked fluorescent proteins in the confocal microscope (**Fig. 5**). We could see that: (i) *P. putida* EM371 recombinants showed brighter fluorescent signal than the EM42 strains, (ii) the fluorescence of cells with attached CFP-AcDoc was dimmer than the fluorescence of their GFP-CtDoc binding counterparts, and (iii) the fluorescence of the EM42 strain expressing the larger AcCoh-CtCoh scaffoldin was more visible than the fluorescence of EM42 cells with displayed single cohesins. The microscopy technique allowed us to also examine positioning of the secreted scaffoldins on the surface of the tested strains. The fluorescence signal was more concentrated in the poles of all EM42 and EM371 recombinants (**Fig. 5**), which indicates asymmetric distribution of the Ag43 autotransporter in the cellular membrane. Such accumulation of overexpressed Ag43 at the polar edges was recently reported also for *E. coli*^69^ and can be a consequence of a cell wall material pushing towards the poles through the continuous lateral insertion of new peptidoglycan building blocks over the rounds of growth and division.^71^

**Figure 5.**
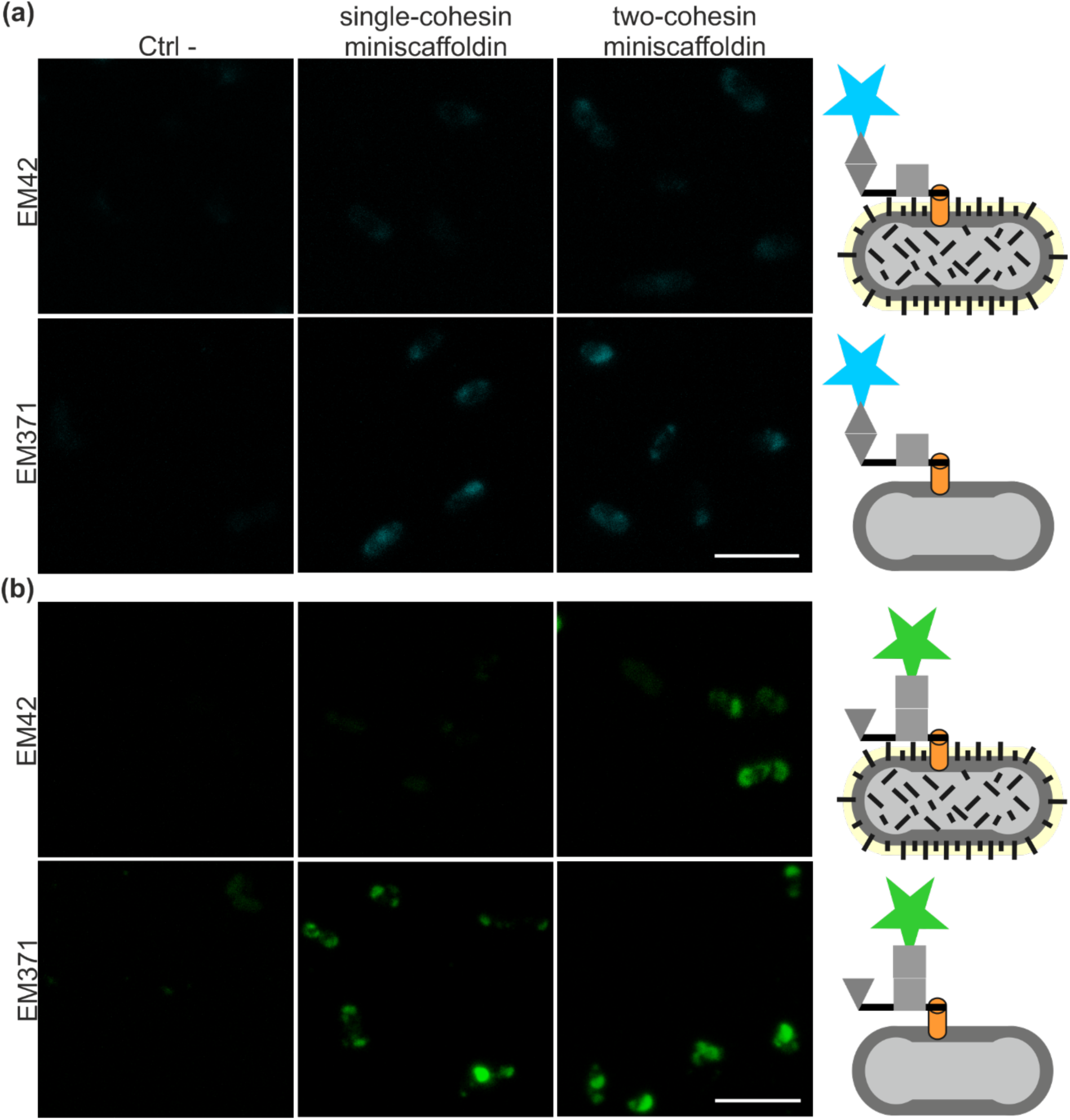
Confocal microscopy of dockerin-tagged fluorescent proteins bound to the *Pseudomonas putida* EM42 and EM371 cells displaying designer scaffoldins *via* the Ag43 autotransporter. **(a)** CFP-AcDoc binding to EM42 or EM371 cells displaying AcCoh or AcCoh-CtCoh scaffoldins. Cells with the pSEVA_*ag43AT* plasmid were used as a negative control (Ctrl-). **(b)** GFP-CtDoc binding to EM42 or EM371 cells displaying CtCoh or AcCoh-CtCoh scaffoldins. All figures are in the same scale, the white bar size is 4 µm.

### Cross-reactivity test and quantification of the scaffoldin molecules displayed on the P. putida surface

Characterized cohesin-dockerin pairs, including those used in this study, are known for their high binding specificity.^24^ However, to rule out any possible cross-reactivity of the binding domains adopted for the aforementioned set of experiments, we added *P. putida* EM371 displaying CtCoh or AcCoh with unmatched BglC-AcDoc or BglC-CtDoc, respectively. The cells showed no β-glucosidase activity upon incubation with pNPG, which confirmed the expected exclusivity of AcCoh-AcDoc and CtCoh-CtDoc bonds (**Fig. S7**). This fact together with the assumed 1:1 dockerin-cohesin binding ratio allowed us to estimate the number of enzyme molecules docked to the surface of a single *P. putida* cell. Knowing the theoretical number of BglC-CtDoc molecules (∼9.49 x 10^12^) in the control reaction with 1 µg of the chimeric enzyme (**Fig. 4b**) and the approximate number of *P. putida* cells (∼1.138 x 10^9^) in suspension used for the whole-cell β-glucosidase activity measurements, we estimated the counts of cohesin-dockerin bonds per bacterium to be ∼1.8 x 10^3^ or ∼4.0 x 10^3^ for the EM42 strain displaying the CtCoh or AcCoh-CtCoh scaffoldins, respectively, and ∼7.8 x 10^3^ or ∼11.0 x 10^3^ for the EM371 strain decorated with CtCoh or AcCoh-CtCoh, respectively. The amount of designer scaffoldins displayed on a single naked *P. putida* EM371 cell *via* an autotransporter system was similar to the number of comparable single- and two-cohesin scaffoldins exported to the surface of the Gram-positive bacterium *L. lactis* in the study of Wieczorek and Martin.^35^ On the other hand, the number of recombinant protein molecules displayed on the EM42 strain did not exceed the values typically reported for Gram-negative hosts such as *E. coli*.^62,72^ These calculations highlighted the superiority of the *P. putida* EM371 over the EM42 strain in terms of scaffoldin secretion efficiency.

### Viability tests and growth of EM42 and EM371 recombinants with surface-docked β-glucosidase in minimal medium with cellobiose

As discussed above, autodisplay of recombinant proteins can affect the viability of a host which might eventually hamper the applicability of a whole-cell biocatalyst. We compared the final optical densities of the cultures conducted to prepare *P. putida* recombinants for the aforementioned assays to evaluate the effect of scaffoldin expression on the viability of EM42 and EM371 strains (**Fig. 6**). None of the cultures showed a substantial drop in OD_600_ after five hours of induction with 3-methylbenzoate when compared with the respective controls. Absorbance of cultures with the EM42 recombinant secreting the two-cohesin scaffoldin was reduced by 9 % (P < 0.05) or 14 % (P < 0.01) when compared with cultures of EM42 pSEVA238_*ag43AT*-*acCoh* or EM42 pSEVA238_*ag43AT-ctCoh*, respectively. However, this minor viability decrease is negligible given the greater capacity of the AcCoh-CtCoh-decorated cells to bind dockerin-tagged proteins (**Figs. 4 and 5**). There were no statistically significant differences in viability among EM371 recombinants with displayed scaffodins (**Fig. 6a**). Only the control strain expressing *ag43AT* alone showed, much to our surprise, ∼20 % lower OD_600_ (P < 0.01) than the remaining EM371 cultures. The same was not seen among EM42 recombinants. Hence we argue that the reduced culture absorbance might be attributed rather to the enhanced clumping of the EM371 pSEVA238_*ag43AT* cells than to their affected viability. The clumping could be caused by the remaining β-helical part of the autotransporter β-domain which also includes an extracellular autochaperone domain. This structure might be more accessible on the surface of the naked strain and promote autoaggregation which is normally associated only with β-helical α domain of the wild-type Ag43.^73^

**Figure 6.**
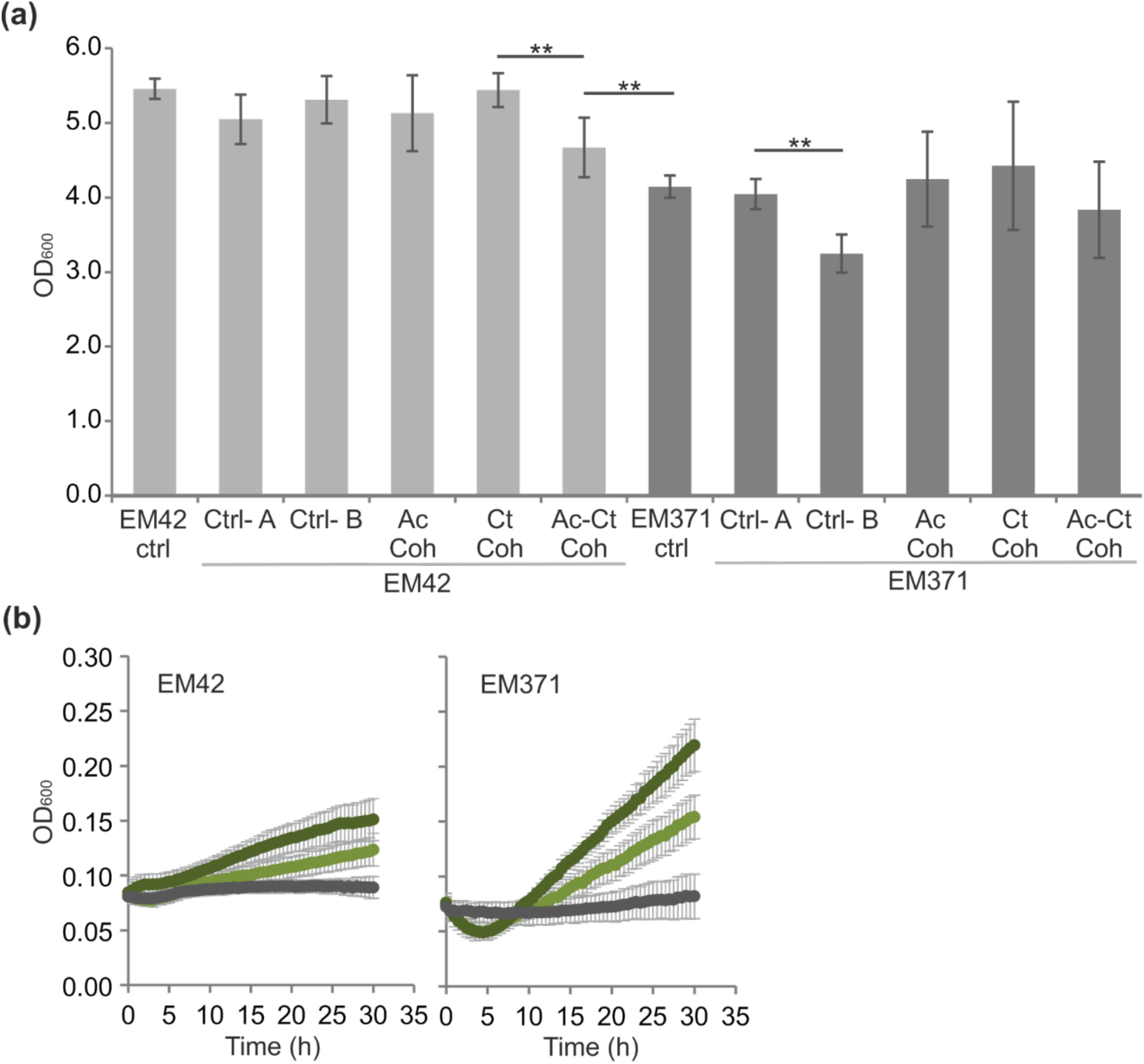
Viability of *Pseudomonas putida* EM42 and EM371 strains displaying designer scaffoldins *via* the Ag43 autotransporter. **(a)** Optical density of *P. putida* cultures measured after 5 h of induction with 0.5 mM 3-methylbenzoate. Used controls were as follows: EM42 ctrl and EM371 ctrl, plasmid-free EM42 and EM371 cells; Ctrl-A, cells with empty pSEVA238; Ctrl-B, cells with pSEVA238_*ag43AT*. Data are shown as mean ± SD from at least three independent experiments, each conducted in two biological replicates. Asterisks denote significance in the difference in between two means at P < 0.01 (**). **(b)** Growth of *P. putida* recombinants with β-glucosidase molecules attached to the displayed two-cohesin miniscaffoldin in minimal medium with cellobiose. EM42 and EM371 cells with displayed AcCoh-CtCoh were incubated with purified BglC-CtDoc only (pale green line) or with a mixture of BglC-CtDoc and BglC-AcDoc (dark green line) at room temperature for 1 h, washed and grown at 30°C in wells of 96-well microtiter plates containing M9 minimal medium with 5 g L^-1^ D-cellobiose used as a sole carbon source. EM42 and EM371 strains bearing pSEVA238b_*ag43AT* plasmid were used as controls (grey line). Error bars show standard deviations from three biological replicates, each conducted in two technical replicates.

It must be emphasized here that all EM371 recombinants and controls grew slower than their EM42 counterparts and their final ODs were ∼20 % lower (P < 0.01) (**Fig. 6a**). This observation is in agreement with the previous study of Martínez-García and co-workers (2020)^46^ and can be attributed to the reduced fitness of the naked strain. Suboptimal viability of EM371, when compared with the robust EM42 chassis, has been a great concern from the beginning of this study. If scaffoldin-bearing *P. putida* strains should serve as reliable enzyme carriers in future real-life projects, they must provide both sufficiently high cell densities and a considerable capacity to attach recombinant proteins to their surface. The data presented in this work, nonetheless, indicate that the benefit of greater display efficiency of EM371 can compensate for lower vigor of this strain. To verify this assumption and to test the two parameters simultaneously, we incubated the AcCoh-CtCoh-displaying EM42 and EM371 strains with either BglC-CtDoc only (we will call here these strains EM42+ and EM371+ for simplification) or with both BglC-CtDoc and BglC-AcDoc (EM42++ and EM371++ in the following text), and we let the washed cells grow in minimal medium with cellobiose used as a sole carbon source (**Fig. 6b**). The assay confirmed that all four tested strains bound such a quantity of β-glucosidase molecules that provided *P. putida* cells with an amount of glucose sufficient for growth. The observed growth was linear most probably due to the constant amount of β-glucosidase molecules in the reaction. Importantly, EM371 recombinants clearly outperformed EM42 strains (**Fig. 6b**). The EM371++ strain grew ∼ 3-times faster than EM42++ (µ = 0.064 h^-1^ and 0.023 h^-1^, respectively) and the EM371+ strain grew ∼ 4-times faster than EM42+ (µ = 0.048 h^-1^ and 0.013 h^-1^, respectively). It is also noteworthy that the EM42++ and EM371++ grew significantly faster (P < 0.01) than the EM42+ and EM371+ strains. This indicated that both cohesins in the AcCoh-CtCoh scaffoldin were occupied by BglC-AcDoc and BglC-CtDoc at the same time, and there was an additive effect of the two BglC molecules. The specific growth rate of EM42+ was ∼57 % of the growth rate of EM42++ and µ of EM371+ was ∼75 % of EM371++’s rate. This result was in agreement with the outcome of the above-described assays (**Fig. 4**) and suggested once again that β-glucosidase was better attached to the cell surface through the CtDoc-CtCoh interaction than through the AcDoc-AcCoh binding pair.

The described experiment was also an addition to our previously published study in which we identified the growth of *P. putida* with cytoplasmic BglC on cellobiose.^4^ Here, we demonstrated that this enzyme can be tagged with dockerin, overproduced in *P. putida*, purified and turned into a cellulosomal mode on the same host, while its activity with a natural substrate is preserved. Intracellular BglC production is nonetheless a more attractive option for our future research. In such a scenario, only two enzymes – an endoglucanase and an exoglucanase or a cellobiohydrolase – need to be assembled in a functional minicellulosome on the surface of a *P. putida* recombinant to enable the decomposition of crystalline cellulose and its utilization for cellular growth. Moreover, well-expressed intracellular β-glucosidase would not become a bottleneck for efficient cellulose hydrolysis as is often reported,^40,74^ but, on the contrary, would secure rapid drain of cellobiose from the cellular surface and prevent inhibition of the remaining cellulases.^75^

## Discussion

Here, we evaluated the capacity of two biotechnologically relevant strains of the Gram-negative bacterium *P. putida*, strains EM42 and EM371 with differences in the complexity of the cellular surface, to display variants of designer scaffoldin proteins and serve as future carriers for cellulosome-like structures.

Single-cohesin and two-cohesin scaffoldins were first prepared together with chimeric dockerin-tagged variants of β-glucosidase BglC and two fluorophores CFP and GFP. These dockerin-tagged reporters allowed direct detection and quantification of displayed scaffoldins in the subsequent assays. A screening among available monomeric autotransporters identified truncated antigen Ag43 from *E. coli* as a potent candidate for display of prepared scaffoldins. The Ag43-based secretion systems have been repeatedly proven useful for display or secretion of recombinant proteins in *E. coli*,^37,38,69^ but, to the best of our knowledge, the use of this autotransporter has not been reported for *P. putida*, thus far.

Single- and two-cohesin scaffoldins displayed *via* the Ag43 autotransporter were then detected on the surfaces of the *P. putida* EM42 and EM371 recombinants. The resulting complexes were not cellulolytic, but their investigation allowed insight into parameters affecting the assembly of such synthetic structures on the surface of a Gram-negative bacterial host. We confirmed that both the nature of a displayed scaffoldin and the character of a host cell’s exterior matter. Out of the two tested cohesin-dockerin pairs, the one from thermophilic *C. thermocellum* showed better bonding in all cases, even though the position of CtCoh in two-cohesin scaffoldin was theoretically less favorable for interaction with the respective dockerin (closer to the cell wall) than that of AcCoh from the mesophilic bacterium *A. cellulolyticus*. Interestingly, the size of the larger two-cohesin scaffoldin appeared not to be a bottleneck for the display. On the contrary, AcCoh-CtCoh was more accessible on the surfaces of the EM42 and EM371 recombinants than the individually displayed cohesins. This is a promising result for our further work, because only parallel assembly of two or more cellulases on the same scaffoldin can promote the substrate channeling effect and result in more efficient whole-cell biocatalysts compared to bacteria displaying or secreting single enzymes.^42,76^

Both two-cohesin and single-cohesin scaffoldins were much more accessible on *P. putida* EM371 than on EM42. We show that up to tens of thousands of scaffoldins can be displayed on the surface of single shaved *P. putida* cell. Nonetheless, the performance of such a strain to be used as a potential biocatalyst must be evaluated from the point of view of both display capacity and the strain’s viability. The EM371 strain with surface-bound β-glucosidase also outperformed the EM42 recombinant in the growth on natural substrate cellobiose. This result indicates that despite its slightly compromised fitness, *P. putida* EM371 shows considerable potential to become a platform for attachment of designer catalytic scaffoldins and that “surface shaving” represents a viable strategy for the design of new Gram-negative bacterial catalysts.

Further improvement of scaffoldin display is possible in both of the discussed *P. putida* strains, because the outer membrane of a Gram-negative bacterium can accommodate up to hundreds of thousands of protein molecules.^77^ More scaffoldins could be available on the cell surface after *e.g.*, optimization of *ag43AT* expression,^52^ removal of periplasmic and outer membrane proteases that can cleave autotransporter or its passenger,^78^ or addition of secretion-promoting CBM to the N-terminus of displayed cohesins.^35^ Besides, the binding of enzymes to the displayed scaffoldins could be further enhanced by the adoption of cohesin-dockerin pairs from extreme thermophiles^22,79^ or by increasing the length of linkers between individual cohesins.^80^ The remaining challenge for *in vivo* assembly of functional designer cellulosomes in a Gram-negative host lies in supplementation of bulky cellulases whose recombinant production and secretion in non-cellulolytic bacteria is problematic.^40,81^ However, as shown in several recent studies, this obstacle can be elegantly bypassed, *e.g*., by the design of a synthetic interkingdom fungal-bacterial consortium^82,83^ or a single-species co-culture of strains with synergistic functions.^84^ These and other strategies are being considered to complement the work described here and pave the way to *P. putida*-based cell factories for the valorization of polymeric waste feedstocks.

## Materials and methods

### Bacterial strains, media and growth conditions

Bacterial strains used in this study are listed in **Ta Supplementary Table S1.** *Escherichia coli* strains employed for cloning or triparental mating and *Pseudomonas putida* strains used for heterologous gene expression were routinely grown in lysogeny broth (LB; 10 g L^-1^ tryptone, 5 g L^-1^ yeast extract, 5 g L^-1^ NaCl, pH 7.0) at 37°C or 30°C, respectively. *P. putida* strains were routinely pre-cultured overnight (15 h) in 2.5 mL of LB medium with agitation of 300 rpm (Heidolph Unimax 1010 and Heidolph Incubator 1000; Heidolph Instruments, Germany). All used solid media (LB or M9 salts minimal medium; per 1 L: 8.5 g Na_2_HPO_4_ 2H_2_O, 3.0 g KH_2_PO_4_, 1.0 g NH_4_Cl, 0.5 g NaCl, pH 7.0) contained 15 g L^-1^ agar. Solid M9 salts media were prepared with 2 mM MgSO_4_, 2.5 mL L^-1^ trace element solution^86^ and 0.2 % (w/v) citrate used as a sole carbon source. Antibiotics of following final concentrations were added to the liquid and solid media to maintain used plasmids: kanamycin (Km) 50 µg mL^-1^, chloramphenicol (Cm) 30 µg mL^-1^, ampicillin (Amp) 150 µg mL^-1^. Growth conditions specific for individual experiments are described in the following sections.

### Plasmid and strain constructions

All used and constructed plasmids from this study are listed in **Supplementary Table S1.** Standard laboratory protocols^87^ were used for DNA manipulations. Oligonucleotide primers used in this study (**Supplementary Table S2**) were purchased from Sigma-Aldrich (USA). Plasmid DNA was isolated with QIAprep Spin Miniprep kit (Qiagen, USA). The genes of interest were amplified by polymerase chain reaction (PCR) using Q5 high fidelity DNA polymerase (New England BioLabs, USA) according to the manufacturer’s protocol. The reaction mixture (50 µL) consisted of polymerase HF or GC buffer (New England BioLabs, USA), water, dNTPs mix (0.2 mM each; Roche, Switzerland), template DNA, primers (0.5 mM each), and DMSO (in case of high GC content in an amplified gene). Two-step overlap extension (OE) PCR^88^ with Q5 DNA polymerase was adopted for the preparation of chimeric genes. PCR products from the first reaction were purified with NucleoSpin Gel and PCR Clean-up (Macherey-Nagel, Germany) and used (1 µL) as templates in the second PCR round. NucleoSpin Gel and PCR Clean-up kit was also routinely used for purification of PCR products either directly from PCR mixture or agarose gels. Colony PCR was performed using NZYTaq II 2x Green Master Mix solution (NZYTech, Portugal). DNA concentration was measured with NanoVue spectrophotometer (GE Healthcare, USA). All restriction enzymes and Quick Ligation kit used for ligation of digested fragments were from New England BioLabs (USA). Digested plasmids and PCR products were separated by DNA electrophoresis with 0.8 % (w/v) agarose gels and visualized using Molecular Imager VersaDoc (Bio-Rad, USA). The flawlessness of PCR-amplified genes cloned into target plasmids was checked by DNA sequencing (Macrogen, South Korea). Chemocompetent *E. coli* cells (CC118 or Dh5α) transformed with plasmids or ligation mixtures were selected on LB agar plates with respective antibiotic, single clones were re-streaked on new LB agar plates with an antibiotic and grown cells were collected in 1 mL of LB with glycerol (20 % w/v) and stored at −80°C. Plasmid constructs were transferred from *E. coli* Dh5α or CC118 donor to *P. putida* EM42 or EM371 by triparental mating, using *E. coli* HB101 helper strain with pRK600 plasmid (**Supplementary Table S1**). Alternatively, electroporation (2.5 kV, 4 – 5 ms pulse) was used for the transformation of *P. putida* cells with selected plasmids using a MicroPulser electroporator and Gene Pulser Cuvettes with 0.2 cm gap (Bio-Rad, USA). Preparation of *P. putida* electrocompetent cells and electroporation procedure itself was performed as described elsewhere.^89^ *P. putida* transconjugants or transformants were selected on M9 agar plates with citrate or on LB agar plates, respectively, with respective antibiotic. All plasmid constructs inserted in *P. putida* were first isolated and re-checked by restriction digestion before the strain was used for further work.

### Preparation of scaffoldin variants, chimaeric proteins, and autotransporters

The original synthetic scaffoldin gene *scaf19L*,^47^ which encodes carbohydrate-binding module CBM3a and cohesin CohCt A2 (named CtCoh in this study) of the cellulosomal-scaffolding protein CipA from *Clostridium thermocellum*, cohesin CohAc C3 (named AcCoh in this study) of the cellulosomal-scaffolding protein ScaC from *Acetivibrio cellulolyticus*, and cohesin CohBc B3 (named BcCoh in this study) of the cellulosomal-scaffolding protein ScaB from *Bacteroides cellulosolvens* interconnected with 27-35 aa long linkers, was PCR amplified using Q5 polymerase and primers Sca19L fw and Sca19L rv (**Supplementary Table S2**) and subcloned into *Nde*I and *Pst*I restriction sites of the modified version of pSEVA238 expression plasmid, pSEVA238b, with synthetic ribosome binding site (RBS).^4^ In parallel, a version of the *scaf19LKT* gene with synthetic RBS was synthesized and codon-optimized for expression in *P. putida* KT2440 (GeneCust, France). The synthetic gene was subcloned from delivery vector pUC57_*scaf19LKT* into *Sac*I and *Kpn*I sites of pSEVA238.

The *bglC* gene encoding β-glucosidase (EC 3.2.1.21) from *Thermobifida fusca*^90^ with N-terminal 6x histidine tag and *ctDoc* gene (encoding *C. thermocellum* dockerin, named here CtDoc, complementary to CtCoh) codon-optimized for expression in *P. putida* KT2440 (GeneCust, France) were separately amplified by PCR from pSEVA238b_*bglC*^4^ and pUC57_*ctDoc*, respectively, using primer pairs BglC-CtDoc TS1F / BglC-CtDoc TS1R and BglC-CtDoc TS2F / BglC-CtDoc TS2R. The first round PCR products were sewed in the second round of OE PCR with TS1F and TS2R primers. The resulting *bglC-ctDoc* chimeric gene was cloned into *Nde*I and *Hind*III sites of pSEVA238b. The pSEVA_*bglC-acDoc* construct bearing *bglC* gene tagged with codon-optimized *A. cellulolyticus* ScaB dockerin, named here AcDoc, was prepared correspondingly using primer pairs BglC-CtDoc TS1F / BglC-AcDoc TS1R and BglC-AcDoc TS2F / BglC-AcDoc TS2R and template plasmids pSEVA238b_*bglC* and pUC57_*acDoc*.

For the purpose of construction of the plasmid allowing the translational fusion of CtDoc to monomeric superfolded GFP (msfGFP), the *gfp* gene was initially amplified with synthetic RBS but without STOP codon from pSEVA238_*gfp* plasmid (SEVA collection) using GFP-N fw and GFP-N rv primers. The PCR product was digested with *Avr*II and *Eco*RI and ligated into pSEVA238, cut with the same pair of enzymes, giving rise to pSEVA238_*gfpN*. The *ctDoc* gene was amplified from pSEVA238_*bglC-acDoc* with its synthetic SGGGS Gly-Ser linker and with its TAA STOP codon using CtDoc fw and CtDoc rv primers. The PCR product was digested with *SacI* and *Hind*III and cloned downstream of the *gfp* gene in pSEVA238_*gfpN*, resulting in pSEVA238_*gfp-ctDoc*. The *gfp-ctDoc* construct was then subcloned into *Nde*I and *HindI*II sites of pET21b for the purpose of gene overexpression and purification of the chimeric protein.

The pSEVA238b_*ag43AT* construct was prepared by subcloning the recombinant Ag43 autotransporter gene from pAg43pol^38^ into *Nde*I and *Hind*III sites of the pSEVA238b vector with synthetic RBS. The *estPAT* gene encoding C-terminal part (331 AA) of EstP esterase autotransporter from *P. putida* KT2440 (PP_0418) with original 23 AA N-terminal leader sequence, E-tag, and polylinker was commercially synthesized (GeneCust, France) and then subcloned from delivery vector pUC57 into *Nde*I and *Hind*III sites of pSEVA238b.

Single cohesin gene *ctCoh* was PCR amplified from pSEVA238_*scaf19LKT* for the purpose of subcloning into the polylinkers of three tested autotransporter systems using primer pair CtCoh fw and CtCoh rv1 (for cloning into *Xho*I and *Bam*HI sites of pSEVA238b_*ag43AT* polylinker) or CtCoh fw and CtCoh rv2 (for cloning into *Eco*RI and *Bam*HI sites of pSEVA238_*igAAT* and pSEVA238_*intAT*). These manipulations gave rise to the constructs pSEVA238_*igAAT-ctCoh*, pSEVA238_*intAT-ctCoh*, and pSEVA238b_*ag43AT-ctCoh*. The pSEVA238b_*ag43AT-acCoh* construct was prepared by subcloning of the *acCoh* gene, PCR amplified from pSEVA238_*scaf19LKT* with primers AcCoh fw and AcCoh rv, into *Xho*I and *Bam*HI sites of pSEVA238b_*ag43AT* polylinker. Similarly, the two-cohesin scaffoldin sequence *acCoh-ctCoh* was amplified from pSEVA238_*scaf19LKT* using the AcCoh-CtCoh fw and CtCoh rv1 primers and inserted into *Xho*I and *Bam*HI sites of pSEVA238b_*ag43AT.*

### Effect of scaffoldin and autotransporter expression on the viability of P. putida cells

Overnight cultures of *P. putida* EM42 cells bearing pSEVA238b_*scaf19L*, pSEVA238_*scaf19LKT*, or empty pSEVA238 were used to inoculate 20 mL of LB with Km in 100 mL shake flasks to initial OD_600_ of 0.05. The cells were grown for three hours with shaking (170 rpm) and then 0.5 mM 3-methylbenzoate (3MB) was added to all cultures to induce expression of recombinant genes. The cells were further cultured for 18 h, and OD_600_ was measured periodically. The effect of IgAAT, IntAT, Ag43AT, and EstPAT autotransporter expression from the pSEVA238 plasmid on viability of the EM42 host was tested correspondingly.

### Analysis of cohesin-dockerin interactions by affinity-based ELISA

Proper folding and ability of AcCoh, CtCoh, and BcCoh cohesins in Scaf19LKT scaffoldin produced in *P. putida* to bind respective dockerins was verified by affinity-based ELISA (enzyme-linked immunosorbent assay). *P. putida* EM42 pSEVA238_*scaf19LKT* was pre-grown overnight in 2.5 mL of LB with Km and 10 mM CaCl_2_. The cells were then used to inoculate 50 mL of LB medium with Km and 10 mM CaCl_2_ in the main culture to the starting OD_600_ of 0.05. The cells were cultured for 3 h at 30°C with shaking (200 rpm), and expression of scaffoldin gene was then induced with 1 mM 3MB. After 5 more hours of growth, the cells were collected by centrifugation (2,500 g, 4 °C, 15 min), washed with ice-cold TBS buffer (25 mM Tris-Cl, 137 mM NaCl, 2.7 mM KCl, pH 7.2) with 10 mM CaCl_2_ and 0.05 % Tween 20, and the cell pellet was frozen at −80 °C overnight. The pellet was then melted and added with 3 mL of TBS buffer, 0.3 mL of PopCulture Reagent (Merck Millipore, USA), 10 µL of DNase I (5 µg mL^-1^; Sigma-Aldrich, USA), and 10 µL of lysozyme (25 mg mL^-1^ stock; Sigma-Aldrich, USA). The cell suspension was incubated for 30 min at room temperature, centrifuged (21,000 g, 4 °C, 30 min), and supernatant (cell-free extract, CFEs) was used for further work. The concentration of total protein in CFE with recombinant Scaf19LKT was determined by Bradford reagent (Sigma-Aldrich, USA), CFE was diluted 1,000-times in 0.1 M sodium carbonate coating buffer (pH 9.0), and 100 µL (the same volume was used for all following steps) were used for overnight coating of individual wells in a MaxiSorp high protein-binding capacity 96-well ELISA plate (Nunc, Denmark) at 4 °C. In the morning, CFE was discarded from the plate, blocking buffer (TBS with 10 mM CaCl_2_ and 0.05 % Tween 20, and 2 % BSA) was added to the wells, and the plate was incubated at room temperature for 1 h (the same incubation conditions were preserved in the remaining steps of the protocol). Blocking buffer was then removed, and the wells were added with fresh blocking buffer containing 0.1 µg ml^-1^ of one of three purified recombinant variants of *Geobacillus sp.* WBI xylanase Xyn-AcDoc, Xyn-CtDoc, or Xyn-BcDoc tagged with a dockerin module from *A. cellulolyticus, C. thermocellum*, or *B. cellulosolvens*, respectively. After the incubation step, the blocking buffer with proteins was discarded, and the wells were washed three times with wash buffer (TBS with 10 mM CaCl_2_ and 0.05 % Tween 20). Anti-Xyn primary antibody diluted 10,000x in the blocking buffer was then added to the wells. After incubation, the washing step was repeated (three washes) and the secondary antibody (HRP-labeled anti-rabbit) diluted 10,000x in blocking buffer was added into the wells. This was followed by incubation and wash steps (four washes) and final detection with 100 µL of 3,3’,5,5’-tetramethylbenzidine (TMB) and Substrate-Chromogen (Dako, Denmark). The reaction occurred for 30 s and then was stopped by the addition of 50 µL of 1 M H_2_SO_4_ per well. The intensity of the resulting color was measured spectrophotometrically at 450 nm.

### Protein purification by affinity chromatography

His-tagged BglC, BglC-AcDoc, BglC-CtDoc, CFP-AcDoc, and GFP-CtDoc were purified from CFE prepared from *P. putida* EM42 or *E. coli* BL21-Gold (DE3) (Agilent Technologies, USA) cells. Overnight cultures in 10 mL of LB with Km and 2 mM CaCl_2_ were inoculated from single colonies of *P. putida* EM42 pSEVA238_*bglC, P. putida* EM42 pSEVA238_*bglC-AcDoc, P. putida* EM42 pSEVA238_*bglC-CtDoc*, or *E. coli* BL21-Gold (DE3) pET28a_*cfp-AcDoc* on LB agar plates. Medium with Amp was used for the *E. coli* BL21-Gold (DE3) pET21b*_gfp-ctDoc* recombinant. Overnight cultures were used for inoculation of 200 mL of fresh LB medium with Km or Amp and 2 mM CaCl_2_ to the final OD_600_ of 0.05. Cells were grown with shaking (170 rpm) for 2.5 h and expression of chimeric genes was induced by 1 mM 3MB (*P. putida* with pSEVA238 plasmids) or 50 µM isopropyl β-D-1-thiogalactopyranoside (*E. coli* with pET plasmids). After induction, the *P. putida* cells were further cultured under the same conditions for another 5 h, while the *E. coli* cultures were grown overnight at a reduced temperature of 20 °C. Cells were then pelleted by centrifugation (2000 g, 15 min, 4 °C), washed with ice-cold purification buffer A (TBS with 10 mM CaCl_2_, 0.05 % Tween 20, and 5 mM imidazole, pH 7.2), centrifuged again, resuspended in 5 mL of the same buffer, and frozen at −80 °C. The next day, the cell suspension was melted, added with ¼ of cOmplete EDTA-free protease inhibitor cocktail tablet (Roche, Switzerland) and 5 µL of Lysonase Bioprocessing Reagent (Merck Millipore, USA) and sonicated 6 x 2 min on ice until the suspension became transparent. The cell lysate was centrifuged (21,000 g, 4 °C, 30 min) and the resulting CFE was collected. Total protein concentration in CFE was determined by Bradford reagent (Sigma-Aldrich, USA), and ∼55 mg of total protein in CFE was incubated for 60 min at 4 °C with 2 mL of Ni-NTA agarose (QIAGEN, Germany) equilibrated with purification buffer A. The slurry was applied to a 10-mL Poly-Prep chromatography column (Bio-Rad, USA), which was then washed with 10 mL of purification buffer B (purification buffer A with 50 mM imidazole). His-tagged protein was eventually eluted with purification buffer C (purification buffer A with 500 mM imidazole) in 1.5 mL fractions. Fractions with the highest β-glucosidase activity or GFP or CFP fluorescence were pooled and applied to an Amicon Ultra centrifugal filter unit with 10 kDa cutoff (Merck Millipore, USA) for protein concentration and buffer exchange (TBS with 10 mM CaCl_2_, 0.05 % Tween 20, and 10 % glycerol). The concentration of purified proteins was determined by Bradford reagent, and proteins were stored at 4 °C or at −20 °C for further use.

### β-glucosidase activity assay

β-glucosidase activity of purified BglC, BglC-CtDoc and BglC-AcDoc was measured using the synthetic substrate *p*-nitrophenyl-*β*-D-glucopyranoside (pNPG; Sigma-Aldrich, USA) following the previously described protocol^4^ with some modifications. Briefly, the reaction mixture (total volume 1,200 µL) contained 1,138 µL of 50 mM sodium phosphate buffer (pH 7.0), 60 µL of pNPG (final conc. 5 mM), and 2 µL of enzyme of concentration adjusted to 0.1 mg mL^-1^. The reaction was run in 1.5 mL test tubes at 37 °C. Samples (600 µL) were withdrawn at 10 and 20 min intervals, mixed with 400 µL of 1 M Na_2_CO_3_ to stop the reaction, and the absorbance of the mixture was measured in a cuvette at 405 nm with a UV/Vis spectrophotometer Ultrospec 2100 (Biochrom, UK). Linearity of the enzymatic reaction during the given time interval was checked prior to these measurements. Specific activity (U mg^-1^) was calculated using the calibration curve, prepared with a *p*-nitrophenol standard (Sigma-Aldrich, USA). One unit (U) of enzymatic activity corresponds to one µmol of *p*-nitrophenol produced per minute.

### Evaluation of scaffoldin display and cohesin-dockerin binding on the P. putida cell surface by β-glucosidase activity assay

The following culture procedure was used for the preparation of cells for subsequent whole-cell assays as well as for evaluation of recombinant protein levels in derived *P. putida* CFEs using SDS-PAGE and western blot analyses. *P. putida* EM42 or EM371 strains, bearing empty pSEVA238 plasmid (negative control) or pSEVA238 plasmid harboring an autotransporter gene with a single- or double-cohesin scaffoldin, were inoculated directly from glycerol stocks into 2.5 mL of LB medium with Km and 10 mM CaCl_2_ and grown overnight with shaking. Overnight cultures were used to inoculate 10 mL of LB medium with Km and 10 mM CaCl_2_ to starting OD_600_ of 0.05. Cells were cultured for 2.5 h with shaking (250 rpm, Unimax 1010 shaker and 1000 Inkubator, Heidolph, Germany), and expression of the autotransporter-scaffoldin chimera was then induced with 0.5 mM 3MB. After 4.5 h of induction, the final OD_600_ was measured, and cells were collected by centrifugation (2000 g, 4 °C, 10 min). Cells were washed with 10 mL of ice-cold TBS buffer with 10 mM CaCl_2_ and 0.05 % Tween 20, centrifuged again and resuspended in 0.5 mL of the same buffer to the OD_600_ of 10.0. Then, 25 µL of purified BglC, BglC-CtDoc or BglC-AcDoc of 1 mg mL^-1^ concentration (∼1.0×10^5^ molecules per cell) were added to the cell suspension, and the whole mixture was incubated in a 1.5 mL test tube for 14 h (overnight) at 4 °C with gentle rotation of the tube (SB2 Rotator, Stuart, UK).

For comparison of the three distinct autotransporters displaying CtCoh on the surface of *P. putida* EM42 or EM371, the cell suspension (250 µL) was washed three times with 1 mL of ice-cold TBS buffer with CaCl_2_ and Tween 20, re-suspended in 250 µL of the same buffer (warmed to room temperature) and added with 5 µL of 100 mM pNPG. The same mixture with EM42 or EM371 cells bearing an empty pSEVA238 plasmid served as a negative control. The mixture in TBS buffer without cells but with purified BglC (1 µL of enzyme of 1 mg mL^-1^ concentration) served as a positive control. The mixtures were incubated for 60 min at 37 °C with shaking (300 rpm, Unimax 1010 shaker and 1000 Inkubator, Heidolph, Germany). Cells were then pelleted by brief centrifugation (2370 g, room temperature, 3 min), supernatants (100 µL) were mixed with 50 µL of 1 M Na_2_CO_3_ in 96-well plate, and absorbance at 405 nm was measured using a Victor^2^ 1420 Multilabel Counter (Perkin Elmer, USA). End-point absorbance values, which corresponded with the activity of the cell surface-bound β-glucosidase molecules, were related to the absorbance measured for free β-glucosidase (positive control) and used for comparison of the displaying capacity of the tested autotransporters in the EM42 and EM371 strains. Possible cross-reactivity of AcDoc with CtCoh and CtDoc with AcCoh was tested using the aforementioned protocol applied on EM371 cells which displayed Ag43AT-CtCoh or Ag43AT-AcCoh fusion and which were incubated with purified BglC-AcDoc or BglC-CtDoc protein, respectively.

Alternatively, for the purpose of quantitative evaluation of BglC-CtDoc or BglC-AcDoc binding to a respective scaffoldin variant (CtCoh, AcCoh, or AcCoh-CtCoh) displayed on EM42 and EM371 cells with Ag43AT, a cell suspension after overnight incubation with complimentary BglC-XDoc chimera was washed three times with ice-cold TBS buffer containing CaCl_2_ and Tween 20, diluted twice with the same buffer (OD_600_ = 5.0), and 0.5 mL was warmed in 1.5 mL test tube for 10 min at 37 °C. Suspension with EM42 or EM371 cells bearing pSEVA238b_*ag43AT* served as negative control. Suspensions were then added with 10 µL of 100 mM pNPG to start the enzymatic reaction. A positive control reaction was run in TBS buffer with the addition of 1 µL of purified BglC-CtDoc or BglC-AcDoc at 1 mg mL^-1^ concentration. Samples (100 µL) were withdrawn every five min (total length of the reaction was 20 min), mixed with 50 µL of 1 M Na_2_CO_3_, and the cells were removed by centrifugation. The supernatant (100 µL) was transferred to the 96-well plate, and absorbance at 405 nm was measured using a Victor^2^ 1420 Multilabel Counter (Perkin Elmer, USA). In the case of low β-glucosidase activity detected, the time intervals of the sample withdrawal were extended to allow accurate activity quantification. Activity (1 U = 1 µM min^-1^) was calculated using a calibration curve prepared with a *p*-nitrophenol standard (Sigma-Aldrich, USA). Activity of the enzyme bound to the surface of each of the tested recombinants was related to the calculated activity of free purified BglC-CtDoc or BglC-AcDoc and used for quantitative comparison of the enzyme-binding capacity of the EM42 and EM371 strains displaying single-cohesin or two-cohesin scaffoldin.

The number of enzyme molecules bound to the surface of a given EM42 or EM371 recombinant was estimated from the molecular weight of BglC-CtDoc (63.46 kDa) or BglC-AcDoc (63.94 kDa) calculated using ExPASy Compute pI/Mw tool (https://web.expasy.org) and from the average number of *P. putida* cells present in 0.5 mL suspension of OD_600_ of 5.0 (∼1.1375×10^9^), determined by counting the cells in samples diluted to OD_600_ of 0.2 in a Bürker chamber (Brand, Germany) using Olympus BX50 microscope with 100x oil immersion objective (Olympus, Japan). Assuming a dockerin:cohesin ratio of 1:1, the calculated amount of β-glucosidase molecules anchored to the cell corresponds to the number of scaffoldins displayed on the cell surface.

### Evaluation of scaffoldin display and cohesion-dockerin binding on the P. putida cell surface using fluorescent proteins

EM42 and EM371 cells for quantitative evaluation of GFP-CtDoc or CFP-AcDoc binding to scaffoldin variants displayed with Ag43AT were prepared as described in the previous section. The cells (0.5 mL, OD_600_=1.0) were incubated overnight at 4 °C with 25 µL of purified dockerin-tagged fluorescent protein of 1 mg mL^-1^ stock concentration (∼1.8×10^5^ fluorophore molecules per cell). After washing and dilution with TBS buffer, the cell suspension was transferred (150 µL per well) to black, clear-bottom, 96-well microplates (Corning, USA), and optical density (600 nm) and GFP (475 nm excitation / 515 nm emission) or CFP (440 nm excitation / 500 nm emission) fluorescence were measured with SpectraMax iD5 microplate reader (Molecular Devices, USA). Recorded fluorescence was normalized by cell density.

### Confocal microscopy

Confocal microscopy was used to visually evaluate GFP-CtDoc or CFP-AcDoc binding to scaffoldin variants displayed with Ag43AT on the surface of EM42 and EM371 strains. Cells bearing pSEVA238b_*ag43AT* were used as a negative control. After incubation with a purified fluorescent protein, a cell suspension was washed and diluted in TBS buffer, and 5 µL were dropped on poly-L-lysine coated glass slides (Sigma-Aldrich, USA) and dried for 60 min at room temperature. Then, the cells were mounted with 5 µL of ProLong antifade reagent (Thermo Fisher Scientific, USA), covered with cover glass, and the slides were analyzed using inverted confocal microscope Leica DMi8 S (Leica Microsystems, Germany) using a 100x oil immersion objective and 6x digital zoom.

### SDS-PAGE and western blot analyses

CFEs and purified proteins for the evaluation of purification procedure were prepared as described in the section on *Protein purification by affinity chromatography*. Cell lysates for the evaluation of expression levels of recombinant proteins in *P. putida* EM42 and EM371 were prepared as follows. Cells, collected from 10 mL of LB medium culture and washed with TBS buffer, were added with 250 uL of B-PER Bacterial Protein Extraction Reagent, 0.25 µL of lysozyme and 0.25 µL of DNase I (all components from Thermo Fisher Scientific, USA) and lysed for 15 min at room temperature with slow agitation. For separation of soluble and insoluble fractions (section *Effect of scaffoldin and autotransporter expression on the viability of P. putida cells.*), cell lysates were centrifuged (21,000 g, 4 °C, 30 min), supernatants (CFE) were collected and pellets (insoluble fractions) were washed with distilled water, centrifuged and re-suspended in 50 µL of distilled water. Samples of CFE, as well as samples of wash, elution, or insoluble fractions or purified chimeric proteins, were added with 5x Laemmli buffer, boiled at 100 °C for 7 min and separated by SDS-PAGE using 12 % gels. In all cases, 5 µg of total protein were loaded per gel well. Gels were stained with Coomassie Brilliant Blue R-250 (Fluka/Sigma-Aldrich, Switzerland).

The staining step was omitted for western blot analyses. Samples of CFE or insoluble fractions (5 µg of total protein per well) were loaded on 12 % polyacrylamide gel. After SDS-PAGE, proteins were electrotransferred during a 30-min interval from the gel onto an Immobilon-P membrane of pore size of 0.45 µm (Merck Millipore, USA) under constant electric current of 0.1 A per gel and voltage of 5-7 V using Trans-Blot SD Semi-Dry Transfer Cell (Bio-Rad, USA). Alternatively, wet transfer was conducted under constant current of 0.375 A per gel using Mini-PROTEAN Tetra Vertical Electrophoresis Cell (Bio-Rad, USA) during a 120-min interval. The membrane was blocked overnight at 4°C in PBS buffer with 3 % (w/v) dry milk and 0.1 % (v/v) Tween 20 and then incubated with mouse anti-6xHis tag monoclonal antibody-HRP conjugate (Clontech or Thermo Fisher Scientific, USA) in the same buffer for 2 h at room temperature. PBS buffer with 1 % (v/v) Tween 20 was employed for membrane washing (4-times 5 min), and the proteins were then visualized with SuperSignal West Pico PLUS Chemiluminescent Substrate (Thermo Fisher Scientific, USA) using FUSION Solo S documentation system (Vilber, Germany).

### Dot blot analysis

*P. putida* recombinants were cultured as described in section *Evaluation of scaffoldin display and cohesin-dockerin binding on P. putida cell surface by β-glucosidase activity assay.* Cells from 10 mL of LB medium culture were collected by centrifugation (2370 g, 4 °C, 5 min), washed twice with ice-cold TBS buffer with 0.05 % Tween 20 (TBS-T) and resuspended in 1 mL of the same buffer to the final OD_600_ of 5.0. Cells were dotted (2 µL per dot) on the nitrocellulose membrane of pore size of 0.45 µm (Thermo Fisher Scientific, USA), and the membrane was dried at 30 °C for 15 min. The membrane was blocked in TBS buffer with 3 % (w/v) dry milk and 0.05 % (v/v) Tween 20 for 30 min at room temperature and then incubated with mouse anti-6xHis tag monoclonal antibody-HRP conjugate (Thermo Fisher Scientific, USA) in TBS-T with 0.05 % BSA for 2 h at room temperature. The membrane was washed 3 times for 5 min with TBS-T and once with TBS, and the His-tagged proteins on the cell surface were then visualized with SuperSignal West Pico PLUS Chemiluminescent Substrate (Thermo Fisher Scientific, USA) using FUSION Solo S documentation system (Vilber, Germany).

### Growth of P. putida recombinants with surface-attached β-glucosidase in minimal medium with cellobiose

Pre-induced *P. putida* EM42 and EM371 cells with displayed AcCoh-CtCoh scaffoldins were prepared as described above. EM42 and EM371 strains, bearing the pSEVA238b_*ag43AT* plasmid, were used as negative controls. The cells (0.5 mL, OD_600_=10.0) were incubated with 25 µL of purified BglC-CtDoc (1 mg mL^-1^) or with a mixture (1:1) of BglC-CtDoc and BglC-AcDoc (25 µL each) at room temperature for 1 h. The suspension was washed three times with ice-cold TBS buffer with CaCl_2_ and Tween 20, and the cells were used to inoculate wells of 96-well microtiter plate containing M9 minimal medium with 2 mM MgSO_4_, 100 µM CaCl_2_, 20 µM FeCl_3_, 2.5 mL L^-1^ trace element solution,^86^ and 5 g L^-1^ D-cellobiose (the only carbon source). The plate with lid was incubated in a Tecan Infinite 200 Pro reader (Tecan, Switzerland) at 30 °C with discontinuous linear shaking, and absorbance at 600 nm was measured at 30 min intervals.

### Statistical analyses

Experiments reported here were conducted at least in two biological replicates (the number of experiments and replicates is specified in figure legends). The mean values and corresponding standard deviations (SD) are presented. When appropriate, data were treated with a two-tailed Student’s t test in Microsoft Office Excel 2013 (Microsoft Corp., USA), and confidence intervals were calculated for a given parameter to manifest a statistically significant difference in means between two experimental datasets.

## Associated Content

Supporting Information

## Notes

The authors declare no competing financial interest.

## Acknowledgements

We thank Prof. Alfredo Martinez from Universidad Nacional Autónoma de México for provided pAg43pol construct, we thank Dr. Sarah Morais and Dr. Johanna Stern from Weizmann Institute of Science, Israel, for valuable discussions and help with the ELISA technique, we thank Dr. Esteban Martínez-García from CNB-CSIC, Spain, for providing *P. putida* EM371 strain and valuable discussions, we also thank Microscopy department of CNB-CSIC for the help with confocal microscopy.

## Funding

This work was funded by the MADONNA (H2020-FET-OPEN-RIA-2017-1-766975), BioRoboost (H2020-NMBP-BIO-CSA-2018), SYNBIO4FLAV (H2020-NMBP/0500) and MIX-UP (H2020-Grant 870294) Contracts of the European Union and the S2017/BMD-3691 InGEMICS-CM Project of the Comunidad de Madrid (European Structural and Investment Funds) as well .well as by the Czech Science Foundation (19-06511Y).

## Notes

### Competing Interest Statement

The authors have declared no competing interest.

## References

(1) Blank, L. M.; Narancic, T.; Mampel, J.; Tiso, T.; O’Connor, K. Biotechnological Upcycling of Plastic Waste and Other Non-Conventional Feedstocks in a Circular Economy. Curr. Opin. Biotechnol. 2020, 62, 212–219. https://doi.org/10.1016/j.copbio.2019.11.011.

(2) Baral, N. R.; Sundstrom, E. R.; Das, L.; Gladden, J.; Eudes, A.; Mortimer, J. C.; Singer, S. W.; Mukhopadhyay, A.; Scown, C. D. Approaches for More Efficient Biological Conversion of Lignocellulosic Feedstocks to Biofuels and Bioproducts. ACS Sustain. Chem. Eng. 2019, 7 (10), 9062–9079. https://doi.org/10.1021/acssuschemeng.9b01229.

(3) Peabody, G. L.; Elmore, J. R.; Martinez-Baird, J.; Guss, A. M. Engineered *Pseudomonas putida* KT2440 Co-Utilizes Galactose and Glucose. Biotechnol. Biofuels 2019, 12 (1), 295. https://doi.org/10.1186/s13068-019-1627-0.

(4) Dvořák, P.; de Lorenzo, V. Refactoring the Upper Sugar Metabolism of *Pseudomonas putida* for Co-Utilization of Cellobiose, Xylose, and Glucose. Metab. Eng. 2018, 48, 94–108. https://doi.org/10.1016/j.ymben.2018.05.019.

(5) Löwe, H.; Sinner, P.; Kremling, A.; Pflüger-Grau, K. Engineering Sucrose Metabolism in *Pseudomonas putida* Highlights the Importance of Porins. Microb. Biotechnol. 2020, 13 (1), 97–106. https://doi.org/10.1111/1751-7915.13283.

(6) Salvachúa, D.; Rydzak, T.; Auwae, R.; De Capite, A.; Black, B. A.; Bouvier, J. T.; Cleveland, N. S.; Elmore, J. R.; Huenemann, J. D.; Katahira, R.; Michener, W. E.; Peterson, D. J.; Rohrer, H.; Vardon, D. R.; Beckham, G. T.; Guss, A. M. Metabolic Engineering of *Pseudomonas putida* for Increased Polyhydroxyalkanoate Production from Lignin. Microb. Biotechnol. 2020, 13 (1), 290–298. https://doi.org/10.1111/1751-7915.13481.

(7) Kohlstedt, M.; Starck, S.; Barton, N.; Stolzenberger, J.; Selzer, M.; Mehlmann, K.; Schneider, R.; Pleissner, D.; Rinkel, J.; Dickschat, J. S.; Venus, J.; B J H van Duuren, J.; Wittmann, C. From Lignin to Nylon: Cascaded Chemical and Biochemical Conversion Using Metabolically Engineered *Pseudomonas putida*. Metab. Eng. 2018, 47, 279–293. https://doi.org/10.1016/j.ymben.2018.03.003.

(8) Franden, M. A.; Jayakody, L. N.; Li, W.-J.; Wagner, N. J.; Cleveland, N. S.; Michener, W. E.; Hauer, B.; Blank, L. M.; Wierckx, N.; Klebensberger, J.; Beckham, G. T. Engineering *Pseudomonas putida* KT2440 for Efficient Ethylene Glycol Utilization. Metab. Eng. 2018, 48, 197–207. https://doi.org/10.1016/j.ymben.2018.06.003.

(9) Molina, L.; Rosa, R. L.; Nogales, J.; Rojo, F. *Pseudomonas putida* KT2440 Metabolism Undergoes Sequential Modifications during Exponential Growth in a Complete Medium as Compounds Are Gradually Consumed. Environ. Microbiol. 2019, 21 (7), 2375–2390. https://doi.org/10.1111/1462-2920.14622.

(10) Kukurugya, M. A.; Mendonca, C. M.; Solhtalab, M.; Wilkes, R. A.; Thannhauser, T. W.; Aristilde, L. Multi-Omics Analysis Unravels a Segregated Metabolic Flux Network That Tunes Co-Utilization of Sugar and Aromatic Carbons in *Pseudomonas putida*. J. Biol. Chem. 2019, jbc.RA119.007885. https://doi.org/10.1074/jbc.RA119.007885.

(11) Kampers, L. F. C.; Volkers, R. J. M.; Martins dos Santos, V. A. P. *Pseudomonas putida* KT2440 Is HV1 Certified, Not GRAS. Microb. Biotechnol. 2019, 12 (5), 845–848. https://doi.org/10.1111/1751-7915.13443.

(12) Nogales, J.; Mueller, J.; Gudmundsson, S.; Canalejo, F. J.; Duque, E.; Monk, J.; Feist, A. M.; Ramos, J. L.; Niu, W.; Palsson, B. O. High-Quality Genome-Scale Metabolic Modelling of *Pseudomonas putida* Highlights Its Broad Metabolic Capabilities. Environ. Microbiol. 2020, 22 (1), 255–269. https://doi.org/10.1111/1462-2920.14843.

(13) Horlamus, F.; Wang, Y.; Steinbach, D.; Vahidinasab, M.; Wittgens, A.; Rosenau, F.; Henkel, M.; Hausmann, R. Potential of Biotechnological Conversion of Lignocellulose Hydrolyzates by *Pseudomonas putida* KT2440 as a Model Organism for a Bio-Based Economy. GCB Bioenergy 2019, 11 (12), 1421–1434. https://doi.org/10.1111/gcbb.12647.

(14) Jayakody, L. N.; Johnson, C. W.; Whitham, J. M.; Giannone, R. J.; Black, B. A.; Cleveland, N. S.; Klingeman, D. M.; Michener, W. E.; Olstad, J. L.; Vardon, D. R.; Brown, R. C.; Brown, S. D.; Hettich, R. L.; Guss, A. M.; Beckham, G. T. Thermochemical Wastewater Valorization via Enhanced Microbial Toxicity Tolerance. Energy Environ. Sci. 2018, 11 (6), 1625–1638. https://doi.org/10.1039/C8EE00460A.

(15) Poblete-Castro, I.; Becker, J.; Dohnt, K.; dos Santos, V. M.; Wittmann, C. Industrial Biotechnology of *Pseudomonas putida* and Related Species. Appl. Microbiol. Biotechnol. 2012, 93 (6), 2279–2290. https://doi.org/10.1007/s00253-012-3928-0.

(16) Sánchez-Pascuala, A.; de Lorenzo, V.; Nikel, P. I. Refactoring the Embden–Meyerhof– Parnas Pathway as a Whole of Portable GlucoBricks for Implantation of Glycolytic Modules in Gram-Negative Bacteria. ACS Synth. Biol. 2017, 6 (5), 793–805. https://doi.org/10.1021/acssynbio.6b00230.

(17) Nikel, P. I.; de Lorenzo, V. *Pseudomonas putida* as a Functional Chassis for Industrial Biocatalysis: From Native Biochemistry to Trans-Metabolism. Metab. Eng. 2018, 50, 142–155. https://doi.org/10.1016/j.ymben.2018.05.005.

(18) Martínez-García, E.; de Lorenzo, V. Molecular Tools and Emerging Strategies for Deep Genetic/Genomic Refactoring of *Pseudomonas*. Curr. Opin. Biotechnol. 2017, 47, 120–132. https://doi.org/10.1016/j.copbio.2017.06.013.

(19) Xu, Q.; Resch, M. G.; Podkaminer, K.; Yang, S.; Baker, J. O.; Donohoe, B. S.; Wilson, C.; Klingeman, D. M.; Olson, D. G.; Decker, S. R.; Giannone, R. J.; Hettich, R. L.; Brown, S. D.; Lynd, L. R.; Bayer, E. A.; Himmel, M. E.; Bomble, Y. J. Dramatic Performance of *Clostridium thermocellum* Explained by Its Wide Range of Cellulase Modalities. Sci. Adv. 2016, 2 (2), e1501254. https://doi.org/10.1126/sciadv.1501254.

(20) Adams, J. J.; Pal, G.; Jia, Z.; Smith, S. P. Mechanism of Bacterial Cell-Surface Attachment Revealed by the Structure of Cellulosomal Type II Cohesin-Dockerin Complex. Proc. Natl. Acad. Sci. 2006, 103 (2), 305–310. https://doi.org/10.1073/pnas.0507109103.

(21) Schoeler, C.; Malinowska, K. H.; Bernardi, R. C.; Milles, L. F.; Jobst, M. A.; Durner, E.; Ott, W.; Fried, D. B.; Bayer, E. A.; Schulten, K.; Gaub, H. E.; Nash, M. A. Ultrastable Cellulosome-Adhesion Complex Tightens under Load. Nat. Commun. 2014, 5 (1), 1–8. https://doi.org/10.1038/ncomms6635.

(22) Gunnoo, M.; Cazade, P.-A.; Bayer, E. A.; Thompson, D. Molecular Simulations Reveal That a Short Helical Loop Regulates Thermal Stability of Type I Cohesin-Dockerin Complexes. Phys. Chem. Chem. Phys. PCCP 2018, 20 (45), 28445–28451. https://doi.org/10.1039/c8cp04800b.

(23) Demain, A. L.; Newcomb, M.; Wu, J. H. D. Cellulase, Clostridia, and Ethanol. Microbiol. Mol. Biol. Rev. MMBR 2005, 69 (1), 124–154. https://doi.org/10.1128/MMBR.69.1.124-154.2005.

(24) Artzi, L.; Bayer, E. A.; Moraïs, S. Cellulosomes: Bacterial Nanomachines for Dismantling Plant Polysaccharides. Nat. Rev. Microbiol. 2017, 15 (2), 83–95. https://doi.org/10.1038/nrmicro.2016.164.

(25) Fierobe, H.-P.; Mingardon, F.; Mechaly, A.; Bélaïch, A.; Rincon, M. T.; Pagès, S.; Lamed, R.; Tardif, C.; Bélaïch, J.-P.; Bayer, E. A. Action of Designer Cellulosomes on Homogeneous Versus Complex Substrates Controlled Incorporation of Three Distinct Enzymes Into a Defined Trifunctional Scaffoldin. J. Biol. Chem. 2005, 280 (16), 16325–16334. https://doi.org/10.1074/jbc.M414449200.

(26) Arfi, Y.; Shamshoum, M.; Rogachev, I.; Peleg, Y.; Bayer, E. A. Integration of Bacterial Lytic Polysaccharide Monooxygenases into Designer Cellulosomes Promotes Enhanced Cellulose Degradation. Proc. Natl. Acad. Sci. U. S. A. 2014, 111 (25), 9109–9114. https://doi.org/10.1073/pnas.1404148111.

(27) You, C.; Zhang, Y.-H. P. Annexation of a High-Activity Enzyme in a Synthetic Three-Enzyme Complex Greatly Decreases the Degree of Substrate Channeling. ACS Synth. Biol. 2014, 3 (6), 380–386. https://doi.org/10.1021/sb4000993.

(28) Tsai, S.-L.; DaSilva, N. A.; Chen, W. Functional Display of Complex Cellulosomes on the Yeast Surface via Adaptive Assembly. ACS Synth. Biol. 2013, 2 (1), 14–21. https://doi.org/10.1021/sb300047u.

(29) Cho, H.-Y.; Yukawa, H.; Inui, M.; Doi, R. H.; Wong, S.-L. Production of Minicellulosomes from *Clostridium cellulovorans* in *Bacillus subtilis* WB800. Appl. Environ. Microbiol. 2004, 70 (9), 5704–5707. https://doi.org/10.1128/AEM.70.9.5704-5707.2004.

(30) Wen, F.; Sun, J.; Zhao, H. Yeast Surface Display of Trifunctional Minicellulosomes for Simultaneous Saccharification and Fermentation of Cellulose to Ethanol. Appl. Environ. Microbiol. 2010, 76 (4), 1251–1260. https://doi.org/10.1128/AEM.01687-09.

(31) Arai, T.; Matsuoka, S.; Cho, H.-Y.; Yukawa, H.; Inui, M.; Wong, S.-L.; Doi, R. H. Synthesis of *Clostridium cellulovorans* Minicellulosomes by Intercellular Complementation. Proc. Natl. Acad. Sci. U. S. A. 2007, 104 (5), 1456–1460. https://doi.org/10.1073/pnas.0610740104.

(32) Stern, J.; Moraïs, S.; Ben-David, Y.; Salama, R.; Shamshoum, M.; Lamed, R.; Shoham, Y.; Bayer, E. A.; Mizrahi, I. Assembly of Synthetic Functional Cellulosomal Structures onto the Cell Surface of *Lactobacillus plantarum*, a Potent Member of the Gut Microbiome. Appl. Environ. Microbiol. 2018, 84 (8). https://doi.org/10.1128/AEM.00282-18.

(33) Fan, L.-H.; Zhang, Z.-J.; Yu, X.-Y.; Xue, Y.-X.; Tan, T.-W. Self-Surface Assembly of Cellulosomes with Two Miniscaffoldins on *Saccharomyces cerevisiae* for Cellulosic Ethanol Production. Proc. Natl. Acad. Sci. U. S. A. 2012, 109 (33), 13260–13265. https://doi.org/10.1073/pnas.1209856109.

(34) Perret, S.; Casalot, L.; Fierobe, H.-P.; Tardif, C.; Sabathe, F.; Belaich, J.-P.; Belaich, A. Production of Heterologous and Chimeric Scaffoldins by *Clostridium acetobutylicum* ATCC 824. J. Bacteriol. 2004, 186 (1), 253–257. https://doi.org/10.1128/jb.186.1.253-257.2004.

(35) Wieczorek, A. S.; Martin, V. J. J. Effects of Synthetic Cohesin-Containing Scaffold Protein Architecture on Binding Dockerin-Enzyme Fusions on the Surface of *Lactococcus lactis*. Microb. Cell Factories 2012, 11, 160. https://doi.org/10.1186/1475-2859-11-160.

(36) Bokinsky, G.; Peralta-Yahya, P. P.; George, A.; Holmes, B. M.; Steen, E. J.; Dietrich, J.; Lee, T. S.; Tullman-Ercek, D.; Voigt, C. A.; Simmons, B. A.; Keasling, J. D. Synthesis of Three Advanced Biofuels from Ionic Liquid-Pretreated Switchgrass Using Engineered *Escherichia coli*. Proc. Natl. Acad. Sci. U. S. A. 2011, 108 (50), 19949–19954. https://doi.org/10.1073/pnas.1106958108.

(37) Wargacki, A. J.; Leonard, E.; Win, M. N.; Regitsky, D. D.; Santos, C. N. S.; Kim, P. B.; Cooper, S. R.; Raisner, R. M.; Herman, A.; Sivitz, A. B.; Lakshmanaswamy, A.; Kashiyama, Y.; Baker, D.; Yoshikuni, Y. An Engineered Microbial Platform for Direct Biofuel Production from Brown Macroalgae. Science 2012, 335 (6066), 308–313. https://doi.org/10.1126/science.1214547.

(38) Muñoz-Gutiérrez, I.; Moss-Acosta, C.; Trujillo-Martinez, B.; Gosset, G.; Martinez, A. Ag43-Mediated Display of a Thermostable β-Glucosidase in *Escherichia coli* and Its Use for Simultaneous Saccharification and Fermentation at High Temperatures. Microb. Cell Factories 2014, 13, 106. https://doi.org/10.1186/s12934-014-0106-3.

(39) Tanaka, T.; Hirata, Y.; Nakano, M.; Kawabata, H.; Kondo, A. Creation of Cellobiose and Xylooligosaccharides-Coutilizing *Escherichia coli* Displaying Both β-Glucosidase and β-Xylosidase on Its Cell Surface. ACS Synth. Biol. 2014, 3 (7), 446–453. https://doi.org/10.1021/sb400070q.

(40) Tozakidis, I. E. P.; Brossette, T.; Lenz, F.; Maas, R. M.; Jose, J. Proof of Concept for the Simplified Breakdown of Cellulose by Combining *Pseudomonas putida* Strains with Surface Displayed Thermophilic Endocellulase, Exocellulase and β-Glucosidase. Microb. Cell Factories 2016, 15 (1), 103. https://doi.org/10.1186/s12934-016-0505-8.

(41) Schulte, M. F.; Tozakidis, I. E. P.; Jose, J. Autotransporter-Based Surface Display of Hemicellulases on *Pseudomonas putida*: Whole-Cell Biocatalysts for the Degradation of Biomass. ChemCatChem 2017, 9 (20), 3955–3964. https://doi.org/10.1002/cctc.201700577.

(42) Moraïs, S.; Shterzer, N.; Lamed, R.; Bayer, E. A.; Mizrahi, I. A Combined Cell-Consortium Approach for Lignocellulose Degradation by Specialized *Lactobacillus plantarum* Cells. Biotechnol. Biofuels 2014, 7, 112. https://doi.org/10.1186/1754-6834-7-112.

(43) Lee, S. Y.; Choi, J. H.; Xu, Z. Microbial Cell-Surface Display. Trends Biotechnol. 2003, 21 (1), 45–52. https://doi.org/10.1016/s0167-7799(02)00006-9.

(44) Martínez-García, E.; Nikel, P. I.; Aparicio, T.; de Lorenzo, V. Pseudomonas 2.0: Genetic Upgrading of *P. putida* KT2440 as an Enhanced Host for Heterologous Gene Expression. Microb. Cell Factories 2014, 13, 159. https://doi.org/10.1186/s12934-014-0159-3.

(45) Lieder, S.; Nikel, P. I.; de Lorenzo, V.; Takors, R. Genome Reduction Boosts Heterologous Gene Expression in *Pseudomonas putida*. Microb. Cell Factories 2015, 14, 23. https://doi.org/10.1186/s12934-015-0207-7.

(46) Martínez-García, E.; Fraile, S.; Rodríguez Espeso, D.; Vecchietti, D.; Bertoni, G.; de Lorenzo, V. The Naked Cell: Emerging Properties of a Surfome-Streamlined *Pseudomonas putida* Strain. 2020, In preparation.

(47) Vazana, Y.; Barak, Y.; Unger, T.; Peleg, Y.; Shamshoum, M.; Ben-Yehezkel, T.; Mazor, Y.; Shapiro, E.; Lamed, R.; Bayer, E. A. A Synthetic Biology Approach for Evaluating the Functional Contribution of Designer Cellulosome Components to Deconstruction of Cellulosic Substrates. Biotechnol. Biofuels 2013, 6 (1), 182. https://doi.org/10.1186/1754-6834-6-182.

(48) Ramos, J. L.; Gonzalez-Carrero, M.; Timmis, K. N. Broad-Host Range Expression Vectors Containing Manipulated Meta-Cleavage Pathway Regulatory Elements of the TOL Plasmid. FEBS Lett. 1988, 226 (2), 241–246. https://doi.org/10.1016/0014-5793(88)81431-5.

(49) Weinel, C.; Nelson, K. E.; Tümmler, B. Global Features of the *Pseudomonas putida* KT2440 Genome Sequence. Environ. Microbiol. 2002, 4 (12), 809–818. https://doi.org/10.1046/j.1462-2920.2002.00331.x.

(50) Grote, A.; Hiller, K.; Scheer, M.; Münch, R.; Nörtemann, B.; Hempel, D. C.; Jahn, D. JCat: A Novel Tool to Adapt Codon Usage of a Target Gene to Its Potential Expression Host. Nucleic Acids Res. 2005, 33 (Web Server issue), W526–W531. https://doi.org/10.1093/nar/gki376.

(51) Meuskens, I.; Michalik, M.; Chauhan, N.; Linke, D.; Leo, J. C. A New Strain Collection for Improved Expression of Outer Membrane Proteins. Front. Cell. Infect. Microbiol. 2017, 7, 464. https://doi.org/10.3389/fcimb.2017.00464.

(52) Jong, W. S. P.; Schillemans, M.; Hagen-Jongman, C. M. ten; Luirink, J.; Ulsen, P. van. Comparing Autotransporter β-Domain Configurations for Their Capacity to Secrete Heterologous Proteins to the Cell Surface. PLOS ONE 2018, 13 (2), e0191622. https://doi.org/10.1371/journal.pone.0191622.

(53) Rutherford, N.; Mourez, M. Surface Display of Proteins by Gram-Negative Bacterial Autotransporters. Microb. Cell Factories 2006, 5, 22. https://doi.org/10.1186/1475-2859-5-22.

(54) Tozakidis, I. E. P.; Sichwart, S.; Jose, J. Going beyond E. Coli: Autotransporter Based Surface Display on Alternative Host Organisms. New Biotechnol. 2015, 32 (6), 644–650. https://doi.org/10.1016/j.nbt.2014.12.008.

(55) Wells, T. J.; Sherlock, O.; Rivas, L.; Mahajan, A.; Beatson, S. A.; Torpdahl, M.; Webb, R. I.; Allsopp, L. P.; Gobius, K. S.; Gally, D. L.; Schembri, M. A. EhaA Is a Novel Autotransporter Protein of Enterohemorrhagic *Escherichia coli* O157:H7 That Contributes to Adhesion and Biofilm Formation. Environ. Microbiol. 2008, 10 (3), 589–604. https://doi.org/10.1111/j.1462-2920.2007.01479.x.

(56) Leščić Ašler, I.; Ivić, N.; Kovačić, F.; Schell, S.; Knorr, J.; Krauss, U.; Wilhelm, S.; Kojić-Prodić, B.; Jaeger, K.-E. Probing Enzyme Promiscuity of SGNH Hydrolases. ChemBioChem 2010, 11 (15), 2158–2167. https://doi.org/10.1002/cbic.201000398.

(57) Klauser, T.; Pohlner, J.; Meyer, T. F. Extracellular Transport of Cholera Toxin B Subunit Using Neisseria IgA Protease Beta-Domain: Conformation-Dependent Outer Membrane Translocation. EMBO J. 1990, 9 (6), 1991–1999.

(58) Freudl, R. Signal Peptides for Recombinant Protein Secretion in Bacterial Expression Systems. Microb. Cell Factories 2018, 17 (1), 52. https://doi.org/10.1186/s12934-018-0901-3.

(59) Veiga, E.; de Lorenzo, V.; Fernández, L. A. Autotransporters as Scaffolds for Novel Bacterial Adhesins: Surface Properties of *Escherichia coli* Cells Displaying Jun/Fos Dimerization Domains. J. Bacteriol. 2003, 185 (18), 5585–5590.

(60) Kjaergaard, K.; Hasman, H.; Schembri, M. A.; Klemm, P. Antigen 43-Mediated Autotransporter Display, a Versatile Bacterial Cell Surface Presentation System. J. Bacteriol. 2002, 184 (15), 4197–4204. https://doi.org/10.1128/jb.184.15.4197-4204.2002.

(61) Wentzel, A.; Christmann, A.; Adams, T.; Kolmar, H. Display of Passenger Proteins on the Surface of *Escherichia coli* K-12 by the Enterohemorrhagic *E. coli* Intimin EaeA. J. Bacteriol. 2001, 183 (24), 7273–7284. https://doi.org/10.1128/JB.183.24.7273-7284.2001.

(62) Salema, V.; Marín, E.; Martínez-Arteaga, R.; Ruano-Gallego, D.; Fraile, S.; Margolles, Y.; Teira, X.; Gutierrez, C.; Bodelón, G.; Fernández, L. Á. Selection of Single Domain Antibodies from Immune Libraries Displayed on the Surface of *E. coli* Cells with Two β-Domains of Opposite Topologies. PloS One 2013, 8 (9), e75126. https://doi.org/10.1371/journal.pone.0075126.

(63) Tozakidis, I. E. P.; Lüken, L. M.; Üffing, A.; Meyers, A.; Jose, J. Improving the Autotransporter-Based Surface Display of Enzymes in *Pseudomonas putida* KT2440. Microb. Biotechnol. 2020, 13 (1), 176–184. https://doi.org/10.1111/1751-7915.13419.

(64) Wagner, S.; Bader, M. L.; Drew, D.; de Gier, J.-W. Rationalizing Membrane Protein Overexpression. Trends Biotechnol. 2006, 24 (8), 364–371. https://doi.org/10.1016/j.tibtech.2006.06.008.

(65) Schlegel, S.; Rujas, E.; Ytterberg, A. J.; Zubarev, R. A.; Luirink, J.; de Gier, J.-W. Optimizing Heterologous Protein Production in the Periplasm of *E. coli* by Regulating Gene Expression Levels. Microb. Cell Factories 2013, 12, 24. https://doi.org/10.1186/1475-2859-12-24.

(66) Van Gerven, N.; Sleutel, M.; Deboeck, F.; de Greve, H.; Hernalsteens, J.-P. Surface Display of the Receptor-Binding Domain of the F17a-G Fimbrial Adhesin through the Autotransporter AIDA-I Leads to Permeability of Bacterial Cells. Microbiol. Read. Engl. 2009, 155 (Pt 2), 468–476. https://doi.org/10.1099/mic.0.022327-0.

(67) Valls, M.; de Lorenzo, V.; Gonzàlez-Duarte, R.; Atrian, S. Engineering Outer-Membrane Proteins in *Pseudomonas putida* for Enhanced Heavy-Metal Bioadsorption. J. Inorg. Biochem. 2000, 79 (1–4), 219–223. https://doi.org/10.1016/s0162-0134(99)00170-1.

(68) van der Woude, M. W.; Henderson, I. R. Regulation and Function of Ag43 (Flu). Annu. Rev. Microbiol. 2008, 62, 153–169. https://doi.org/10.1146/annurev.micro.62.081307.162938.

(69) Ahan, R. E.; Kırpat, B. M.; Saltepe, B.; Şeker, U. Ö. Ş. A Self-Actuated Cellular Protein Delivery Machine. ACS Synth. Biol. 2019, 8 (4), 686–696. https://doi.org/10.1021/acssynbio.9b00062.

(70) Shaner, N. C.; Steinbach, P. A.; Tsien, R. Y. A Guide to Choosing Fluorescent Proteins. Nat. Methods 2005, 2 (12), 905–909. https://doi.org/10.1038/nmeth819.

(71) Laloux, G.; Jacobs-Wagner, C. How Do Bacteria Localize Proteins to the Cell Pole? J. Cell Sci. 2014, 127 (1), 11–19. https://doi.org/10.1242/jcs.138628.

(72) Hörnström, D.; Larsson, G.; van Maris, A. J. A.; Gustavsson, M. Molecular Optimization of Autotransporter-Based Tyrosinase Surface Display. Biochim. Biophys. Acta BBA - Biomembr. 2019, 1861 (2), 486–494. https://doi.org/10.1016/j.bbamem.2018.11.012.

(73) Klemm, P.; Hjerrild, L.; Gjermansen, M.; Schembri, M. A. Structure-Function Analysis of the Self-Recognizing Antigen 43 Autotransporter Protein from *Escherichia coli*. Mol. Microbiol. 2004, 51 (1), 283–296. https://doi.org/10.1046/j.1365-2958.2003.03833.x.

(74) Sørensen, A.; Lübeck, M.; Lübeck, P. S.; Ahring, B. K. Fungal Beta-Glucosidases: A Bottleneck in Industrial Use of Lignocellulosic Materials. Biomolecules 2013, 3 (3), 612–631. https://doi.org/10.3390/biom3030612.

(75) Gefen, G.; Anbar, M.; Morag, E.; Lamed, R.; Bayer, E. A. Enhanced Cellulose Degradation by Targeted Integration of a Cohesin-Fused β-Glucosidase into the *Clostridium thermocellum* Cellulosome. Proc. Natl. Acad. Sci. U. S. A. 2012, 109 (26), 10298–10303. https://doi.org/10.1073/pnas.1202747109.

(76) Zhang, Y.-H. P. Substrate Channeling and Enzyme Complexes for Biotechnological Applications. Biotechnol. Adv. 2011, 29 (6), 715–725. https://doi.org/10.1016/j.biotechadv.2011.05.020.

(77) Smit, J.; Kamio, Y.; Nikaido, H. Outer Membrane of Salmonella Typhimurium: Chemical Analysis and Freeze-Fracture Studies with Lipopolysaccharide Mutants. J. Bacteriol. 1975, 124 (2), 942–958.

(78) Jose, J.; Bernhardt, R.; Hannemann, F. Cellular Surface Display of Dimeric Adx and Whole Cell P450-Mediated Steroid Synthesis on *E. coli*. J. Biotechnol. 2002, 95 (3), 257–268. https://doi.org/10.1016/s0168-1656(02)00030-5.

(79) Kahn, A.; Moraïs, S.; Galanopoulou, A. P.; Chung, D.; Sarai, N. S.; Hengge, N.; Hatzinikolaou, D. G.; Himmel, M. E.; Bomble, Y. J.; Bayer, E. A. Creation of a Functional Hyperthermostable Designer Cellulosome. Biotechnol. Biofuels 2019, 12 (1), 44. https://doi.org/10.1186/s13068-019-1386-y.

(80) Hammel, M.; Fierobe, H.-P.; Czjzek, M.; Kurkal, V.; Smith, J. C.; Bayer, E. A.; Finet, S.; Receveur-Bréchot, V. Structural Basis of Cellulosome Efficiency Explored by Small Angle X-Ray Scattering. J. Biol. Chem. 2005, 280 (46), 38562–38568. https://doi.org/10.1074/jbc.M503168200.

(81) Becker, S.; Theile, S.; Heppeler, N.; Michalczyk, A.; Wentzel, A.; Wilhelm, S.; Jaeger, K.-E.; Kolmar, H. A Generic System for the *Escherichia coli* Cell-Surface Display of Lipolytic Enzymes. FEBS Lett. 2005, 579 (5), 1177–1182. https://doi.org/10.1016/j.febslet.2004.12.087.

(82) Minty, J. J.; Singer, M. E.; Scholz, S. A.; Bae, C.-H.; Ahn, J.-H.; Foster, C. E.; Liao, J. C.; Lin, X. N. Design and Characterization of Synthetic Fungal-Bacterial Consortia for Direct Production of Isobutanol from Cellulosic Biomass. Proc. Natl. Acad. Sci. 2013, 110 (36), 14592–14597. https://doi.org/10.1073/pnas.1218447110.

(83) Zhang, S.; Merino, N.; Okamoto, A.; Gedalanga, P. Interkingdom Microbial Consortia Mechanisms to Guide Biotechnological Applications. Microb. Biotechnol. 2018, 11 (5), 833–847. https://doi.org/10.1111/1751-7915.13300.

(84) Honjo, H.; Iwasaki, K.; Soma, Y.; Tsuruno, K.; Hamada, H.; Hanai, T. Synthetic Microbial Consortium with Specific Roles Designated by Genetic Circuits for Cooperative Chemical Production. Metab. Eng. 2019, 55, 268–275. https://doi.org/10.1016/j.ymben.2019.08.007.

(85) Yang, J.; Zhang, Y. I-TASSER Server: New Development for Protein Structure and Function Predictions. Nucleic Acids Res. 2015, 43 (Web Server issue), W174–W181. https://doi.org/10.1093/nar/gkv342.

(86) Abril, M. A.; Michan, C.; Timmis, K. N.; Ramos, J. L. Regulator and Enzyme Specificities of the TOL Plasmid-Encoded Upper Pathway for Degradation of Aromatic Hydrocarbons and Expansion of the Substrate Range of the Pathway. J. Bacteriol. 1989, 171 (12), 6782–6790. https://doi.org/10.1128/jb.171.12.6782-6790.1989.

(87) Sambrook, J.; Russell, D. W. Molecular Cloning: A Laboratory Manual; CSHL Press, 2001.

(88) Horton, R. M.; Cai, Z.; Ho, S. N.; Pease, L. R. Gene Splicing by Overlap Extension: Tailor-Made Genes Using the Polymerase Chain Reaction. BioTechniques 1990, 8 (5), 528–535.

(89) Aparicio, T.; de Lorenzo, V.; Martínez-García, E. Broadening the SEVA Plasmid Repertoire to Facilitate Genomic Editing of Gram-Negative Bacteria. In Hydrocarbon and Lipid Microbiology Protocols: Genetic, Genomic and System Analyses of Pure Cultures; McGenity, T. J., Timmis, K. N., Nogales, B., Eds.; Springer Protocols Handbooks; Springer: Berlin, Heidelberg, 2017; pp 9–27. https://doi.org/10.1007/8623_2015_102.

(90) Spiridonov, N. A.; Wilson, D. B. Cloning and Biochemical Characterization of BglC, a β-Glucosidase from the Cellulolytic Actinomycete *Thermobifida fusca*. Curr. Microbiol. 2001, 42 (4), 295–301. https://doi.org/10.1007/s002840110220.

