## Supplemental information for "Surface display of designer protein scaffolds on genome-reduced strains of *Pseudomonas putida*"

by

Pavel Dvořák[1*], Edward A. Bayer[2], and Víctor de Lorenzo[3*]

[1]*Department of Experimental Biology (Microbiology Section), Faculty of Science, Masaryk University, Kamenice 753/5, 62500, Brno, Czech Republic.*

[2] *Department of Biomolecular Sciences, The Weizmann Institute of Science, Rehovot 76100, Israel*

[3]*Systems and Synthetic Biology Program, Centro Nacional de Biotecnología CNB-CSIC, Cantoblanco, Darwin 3, 28049 Madrid, Spain.*

**Keywords:** *Pseudomonas putida*, cellulosome, designer scaffoldin, surface display, synthetic biology

* Co-corresponding authors:

Prof. V. de Lorenzo

Systems and Synthetic Biology Program, Centro Nacional de Biotecnología (CNB-CSIC)

Darwin 3, Campus de Cantoblanco Madrid 28049, Spain

Dr. Pavel Dvořák

Department of Experimental Biology (Section of Microbiology), Faculty of Science

Masaryk University, Kamenice 735/5, Brno 62500, Czech Republic

### Contents

**Supplementary tables** ……………………………………………………………………3

**Table S1.** Strains and plasmids used in this study.

**Table S2.** Oligonucleotide primers used in this study.

**Supplementary figures** ………………………………………………...………………7

**Figure S1.** Schematic reconstruction of genomic deletions in *Pseudomonas putida* strains EM42 and EM371 used in this study.

**Figure S2.** Effect of codon optimization of the *scaf19L* gene on toxicity, solubility, and stability of the corresponding protein produced in *Pseudomonas putida* EM42.

**Figure S3.** SDS-polyacrylamide gels with samples from purification of BglC, BglC-CtDoc, Bglc-AcDoc, GFP-CtDoc, and CFP-AcDoc.

**Figure S4.** Effect of an autotransporter expression on viability of the host bacterium *Pseudomonas putida* EM42.

**Figure S5.** Expression and display analysis of three different autotransporters with CtCoh passenger in *Pseudomonas putida* EM42 and EM371.

**Figure S6.** Expression analysis of three miniscaffoldin variants displayed *via* Ag43 autotransporter in *Pseudomonas putida* EM42 and EM371.

**Figure S7.** Cross-reactivity test with used cohesin-dockerin pairs.

**Supplementary sequences** ……………………………………………………………….15

**References**………………………………………………………...…………………19

### Supplementary tables

**Table S1.** Strains and plasmids used in this study.

| Strain or plasmid | Characteristics | Source or reference |
|---|---|---|
| ***Escherichia coli*** | | |
| Dh5α | Cloning host: F-λ-*endA1 glnX44*(AS) *thiE1 recA1 relA1 spoT1 gyrA96*(NalR) *rfbC1 deoR nupG* Φ80(*lacZ*Δ*M15*) Δ(*argF-lac*)*U169 hsdR17*($r_K^- m_K^+$) | Grant *et al.*[1] |
| CC118 | *Cloning host:* Δ*(ara-leu) araD* Δ*lac X174 galE galK phoA thiE1 rpoB(Rif*$^R$*) argE(Am) recA1* | Manoil and Beckwith[2] |
| BL21-Gold (DE3) | Expression host, *E. coli* B derivative: $F^-$ *ompT hsdS*($r_B^-$ $m_B^-$) *dcm*$^+$ *Tet*$^r$ *gal* λ(DE3 [*lacI*$^Q$ *lacUV5-T7 gene 1 ind1 sam7 nin5*]) *endA* Hte | Agilent Technologies, USA |
| HB101 | Helper strain for tri-parental mating: F– λ– *hsdS20*($r_B^-$ $m_B^-$) *recA13 leuB6*(Am) *araC14* Δ(*gpt-proA*)62 *lacY1 galK2*(OC) *xyl-5 mtl-1 thiE1 rpsL20*(Sm$^R$) *glnX44*(AS) | Boyer and Roulland-Dussoix[3] |
| ***Pseudomonas putida*** | | |
| EM42 | Derivative of strain *P. putida* KT2440: Δprophages1,2,3,4 ΔTn7 Δ*endA1* Δ*endA2* Δ*hsdRMS* Δflagellum ΔTn*4652* | Martínez-García *et al.*[4] |
| EM371 | Derivative of strain *P. putida* KT2440: Δprophage4 ΔTn7 Δflagellum Δpili Δcurli Δmotility proteins Δalginate biosynthesis Δtwitching motility protein Δsurface adhesion protein Δcellulose synthesis Δouter membrane lipoprotein Δglycosyl transferase | Martínez-García *et al.*[5] |
| **plasmids** | | |
| pRK600 | Helper plasmid for tri-parental mating: *oriV*(*ColE1*) RK2 *tra*$^+$ *mob*$^+$, Cm$^R$ | Kessler *et al.*[6] |
| pSEVA238 | Expression vector: *oriV*(pBBR1) *xylS-Pm neo*, Km$^R$ | Silva-Rocha *et al.*[7] |
| pSEVA238b | Derivative of pSEVA238 with synthetic RBS and adjacent *Nde*I site for subcloning of genes to be expressed | Dvořák and de Lorenzo[8] |
| pSEVA238_*gfp* | pSEVA238 bearing synthetic RBS and gene of monomeric superfolder GFP (msfGFP) cloned into *Hind*III and *Spe*I sites of SEVA polylinker | SEVA collection |
| pET21b | Expression vector: *ori*(pBR322, f1), T7 promoter and terminator, *lacI*, Amp$^R$ | Merck, USA |
| pET28a_*scaf19L* | Expression vector: *ori*(pBR322, f1), T7 promoter and terminator, *lacI*, Km$^R$, with synthetic scaffoldin gene *scaf19L* | Vazana *et al.*[9] |

| | | |
|---|---|---|
| | (encodes CBM3a and CohCt A2 of CipA from *Clostridium thermocellum*, CohAc C3 of ScaC from *Acetivibrio cellulolyticus*, CohBc B3 of ScaB from *Bacteroides cellulosolvens* with 27-35 aa long linkers, and C-terminal 6xHis tag) cloned into *Nco*I and *Xho*I sites | |
| pSEVA238b_*scaf19L* | pSEVA238b with scaffoldin gene *scaf19L* cloned into *Nde*I and *Pst*I sites | This study |
| pUC57_*scaf19LKT* | Cloning vector (Amp$^R$) with *scaf19L* gene synthesized with RBS and codon-optimized for expression in *P. putida* KT2440 | This study |
| pSEVA238_*scaf19LKT* | pSEVA238 with synthesized *scaf19L* gene (with RBS) cloned into *Sac*I and *Kpn*I sites | This study |
| pUC57_*ctDoc* | Cloning vector (Amp$^R$) with synthetic *ctDoc* sequence (encodes *C. thermocellum* dockerin with C-terminal 6xHis tag) codon-optimized for expression in *P. putida* KT2440 | This study (GeneCust, France) |
| pUC57_*acDoc* | Cloning vector (Amp$^R$) with synthetic *acDoc* sequence (encodes *A. cellulolyticus* dockerin with C-terminal 6xHis tag) codon-optimized for expression in *P. putida* KT2440 | This study (GeneCust, France) |
| pSEVA238b_*bglC* | pSEVA238b harboring *bglC* gene of β-glucosidase from *Thermobifida fusca* with N-terminal 6xHis tag cloned into *Nde*I and *Hind*III sites | Dvořák and de Lorenzo[8] |
| pSEVA238b_*bglC-ctDoc* | pSEVA238b harboring chimeric *bglC-ctDoc* gene cloned into *Nde*I and *Hind*III sites | This study |
| pSEVA238b_*bglC-acDoc* | pSEVA238b harboring chimeric *bglC-acDoc* gene cloned into *Nde*I and *Hind*III sites | This study |
| pSEVA238_*gfpN* | pSEVA238 with *gfp* gene encoding monomeric superfolder green fluorescent protein (msfGFP) with synthetic RBS but lacking STOP codon (*Avr*II/*Eco*RI) | This study |
| pSEVA238_*gfp-ctDoc* | pSEVA238_*gfpN* with *ctDoc* gene cloned in frame downstream the *gfp* gene (*Sac*I/*Hind*III) to form a translational fusion | This study |
| pET21b_*gfp-ctDoc* | pET21b with *gfpN-ctDoc* cloned into *Nde*I and *Hind*III sites. | This study |
| pET28a_*cfp-acDoc* | pET28a with *cfp-acDoc* gene (encodes cyan fluorescent protein mCerulean fused to *A. cellulolyticus* ScaB dockerin) cloned into *Xba*I and *Xho*I sites | Gift from Edward A. Bayer |
| pUC57_*estPAT* | Cloning vector (Amp$^R$) with synthetic *estPAT* gene encoding C-terminal part (331 AA) of EstP esterase autotransporter from *P. putida* KT2440 with original 23 AA N-terminal leader sequence and polylinker | This study (GeneCust, France) |
| pAg43pol | pTrc99A2 derivative (*trc* promoter) for recombinant protein display using *E. coli* autotransporter Ag43, Amp$^R$ | Muñoz-Gutiérrez *et al.*[10] |

| | | |
|---|---|---|
| pSEVA238_*igAAT* | pSEVA238 bearing synthetic RBS and gene encoding C-terminal part (409 AA) of IgA-specific serine endopeptidase autotransporter from *Neisseria gonorrhoeae* with 22 AA N-terminal PelB leader sequence of pectate lyase B of *Erwinia carotovora* CE and SEVA polylinker, whole gene is cloned into *Avr*II and *Spe*I sites | Esteban Martínez-García |
| pSEVA238_*intAT* | pSEVA238 bearing synthetic RBS and gene encoding N-terminal part (654 AA) of intimin γ from enterohemorrhagic *E. coli* O157:H7 with original 39 AA leader sequence, SEVA polylinker is at 3' terminus, whole gene is cloned into *Avr*II and *Spe*I sites | Esteban Martínez-García |
| pSEVA238b_*ag43AT* | pSEVA238b with gene encoding C-terminal part (487 AA) of adhesin Ag43 autotransporter from *E. col* with original 52 AA leader sequence and inserted polylinker, whole gene is cloned into *Nde*I and *Hind*III sites | This study |
| pSEVA238b_*estPAT* | pSEVA238b with gene encoding *estPAT* autotransporter cloned into *Nde*I and *Hind*III sites | This study |
| pSEVA238_*igAAT-ctCoh* | pSEVA238_*igAAT* with gene encoding *ctCoh* cohesin codon-optimized for expression in *P. putida* KT2440 and cloned into *Eco*RI and *Bam*HI sites of *igAAT* polylinker | This study |
| pSEVA238_*intAT-ctCoh* | pSEVA238_*intAT* with gene encoding *ctCoh* cohesin codon-optimized for expression in *P. putida* KT2440 and cloned into *Eco*RI and *Bam*HI sites of *intAT* polylinker | This study |
| pSEVA238b_*ag43AT-ctCoh* | pSEVA238b_*ag43AT* with gene encoding *ctCoh* cohesin codon-optimized for expression in *P. putida* KT2440 and cloned into *Xho*I and *Bam*HI sites of *Ag43AT* polylinker | This study |
| pSEVA238b_*ag43AT-acCoh* | pSEVA238b_*ag43AT* with gene encoding *acCoh* cohesin codon-optimized for expression in *P. putida* KT2440 and cloned into *Xho*I and *Bam*HI sites of *Ag43AT* polylinker | This study |
| pSEVA238b_*ag43AT-acCoh-ctCoh* | pSEVA238b_*ag43AT* with gene encoding *acCoh- ctCoh* scaffoldin codon-optimized for expression in *P. putida* KT2440 and cloned into *Xho*I and *Bam*HI sites of *Ag43AT* polylinker | This study |

Abbreviations: RBS, ribosome binding site; CBM, carbohydrate binding module; Cm, chloramphenicol; Amp, ampicillin; Km, kanamycin; Sm, streptomycin; Sp, spektinomycin; Coh, cohesin; Doc, dockerin.

**Table S2.** Oligonucleotide primers used in this study.

| Name | Sequence (5’→3’)[a] |
|---|---|
| Sca19L fw (*Nde*I) | taacatatggcaaatacaccggtatcag |
| Sca19L rv (*Pst*I) | aatctgcagtcagtggtggtggtggtg |
| BglC-CtDoc TS1F (*Nde*I) | atacatatgcaccatcaccatcac |
| BglC-CtDoc TS1R | ggaaccacccccggattcctgtccgaagattccc |
| BglC-CtDoc TS2F | tccgggggtggttcc |
| BglC-CtDoc TS2R (*Hind*III) | attaagcttttagtggtggtgatggtggtggttcttgtacggcagcgtgtc |
| BglC-AcDoc TS1R | cacgtcgccgtagatgaacttgctagcggaaccacccc |
| BglC-AcDoc TS2F | aagttcatctacggcgacgtg |
| BglC-AcDoc TS2R *(Hind*III) | taaaagcttttagtgatggtgatggtggtg |
| CtCoh fw (*Eco*RI, *Xho*I) | atagaattcagtcgacctcgagtccgatggcgtagtagtgg |
| CtCoh rv1 (*Bam*HI) | tatctgcagggatccgtgatggtggtgatggtg ggtggcgttgccc |
| CtCoh rv2 (*Bam*HI) | tatggatccatgtgatggtggtgatggtgggtggcgttgccc |
| AcCoh fw (*Xho*I) | atactcgagggttccgatttgcaggtggac |
| AcCoh rv (*Bam*HI) | tatggatccgtgatggtggtgatggtggctggcgatcacctcgatc |
| AcCoh-CtCoh fw (*Xho*I) | atagaattcagtcgacctcgagggttccgatttgcaggtg |
| GFP-N fw (*Avr*II) | attcctaggtagcaagaggaatatcatatgagtaaaggagaagaacttttcac |
| GFP-N rv (*Eco*RI) | aatgaattctttgtatagttcatccatgcc |
| CtDoc fw (*Sac*I) | aatgagctctccgggggtggttcc |
| CtDoc rv (*Hind*III) | cgcaagcttttagtggtggtg |

[a] Restriction sites are underlined.

### Supplementary figures

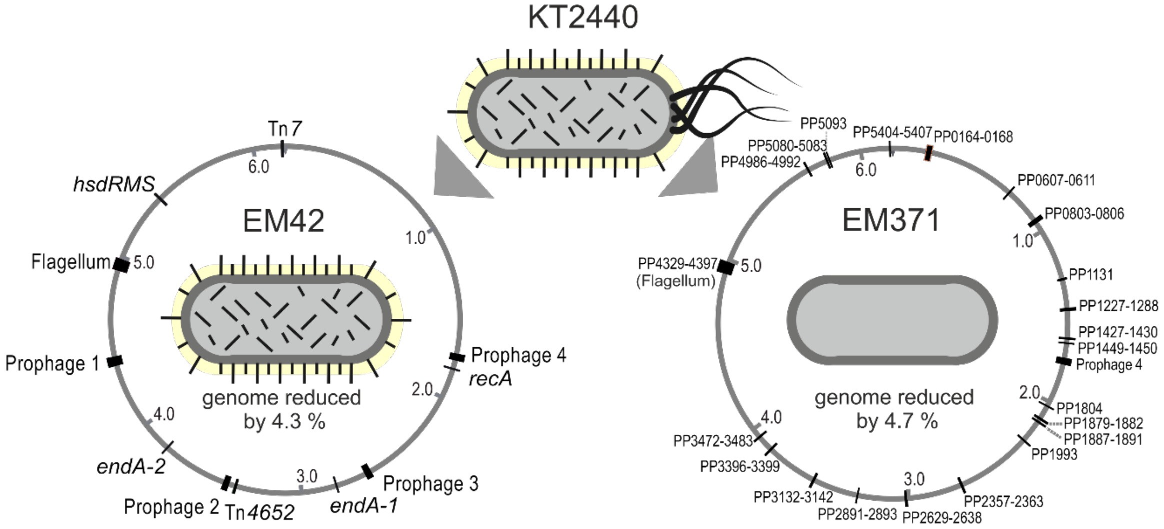


**Figure S1. Schematic reconstruction of genomic deletions in *Pseudomonas putida* strains EM42 and EM371 used in this study.** For more details on *P. putida* EM42 and EM371 see original studies by Martínez-García *et al.*[4] and Martínez-García *et al.*[5], respectively.

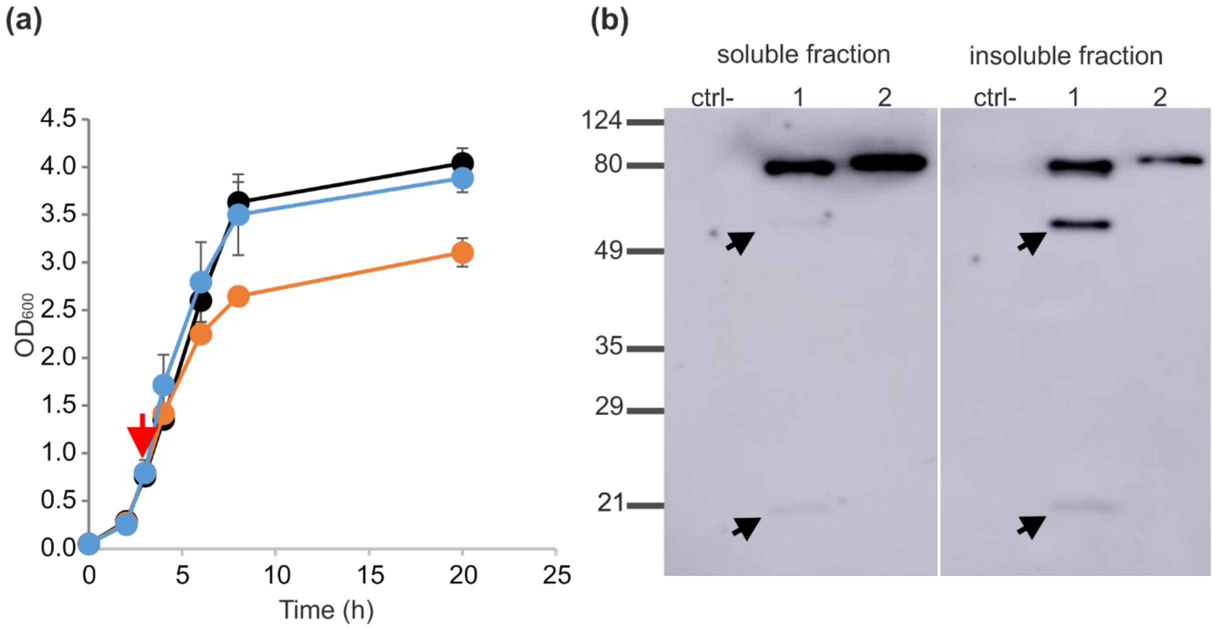


**Figure S2. Effect of codon optimization of the *scaf19L* gene on toxicity, solubility, and stability of the corresponding protein produced in *Pseudomonas putida* EM42. (a)** Effect of *scaf19L* (orange line) and s*caf19LKT* (blue line) expression on viability of *P. putida* EM42 host. Expression of the heterologous gene cloned in pSEVA238 plasmid was induced 3 h after culture start (red arrow) with 0.5 mM 3-methylbenzoate. *P. putida* EM42 with empty pSEVA238 plasmid (black line) was used as a control. Data are shown as mean ± SD from two biological replicates. **(b)** Western blot analysis of Scaf19L (1) and Scaf19LKT (2) with C-terminal 6xHis tag in soluble (cell-free extract) and insoluble fraction of cell lysate. Protein of the theoretical molecular weight of 75.4 kDa was detected using mouse anti-6xHis tag monoclonal antibody-HRP conjugate (Clontech, USA). Black arrows indicate possible products of Scaf19L proteolysis. Protein marker is in kDa.

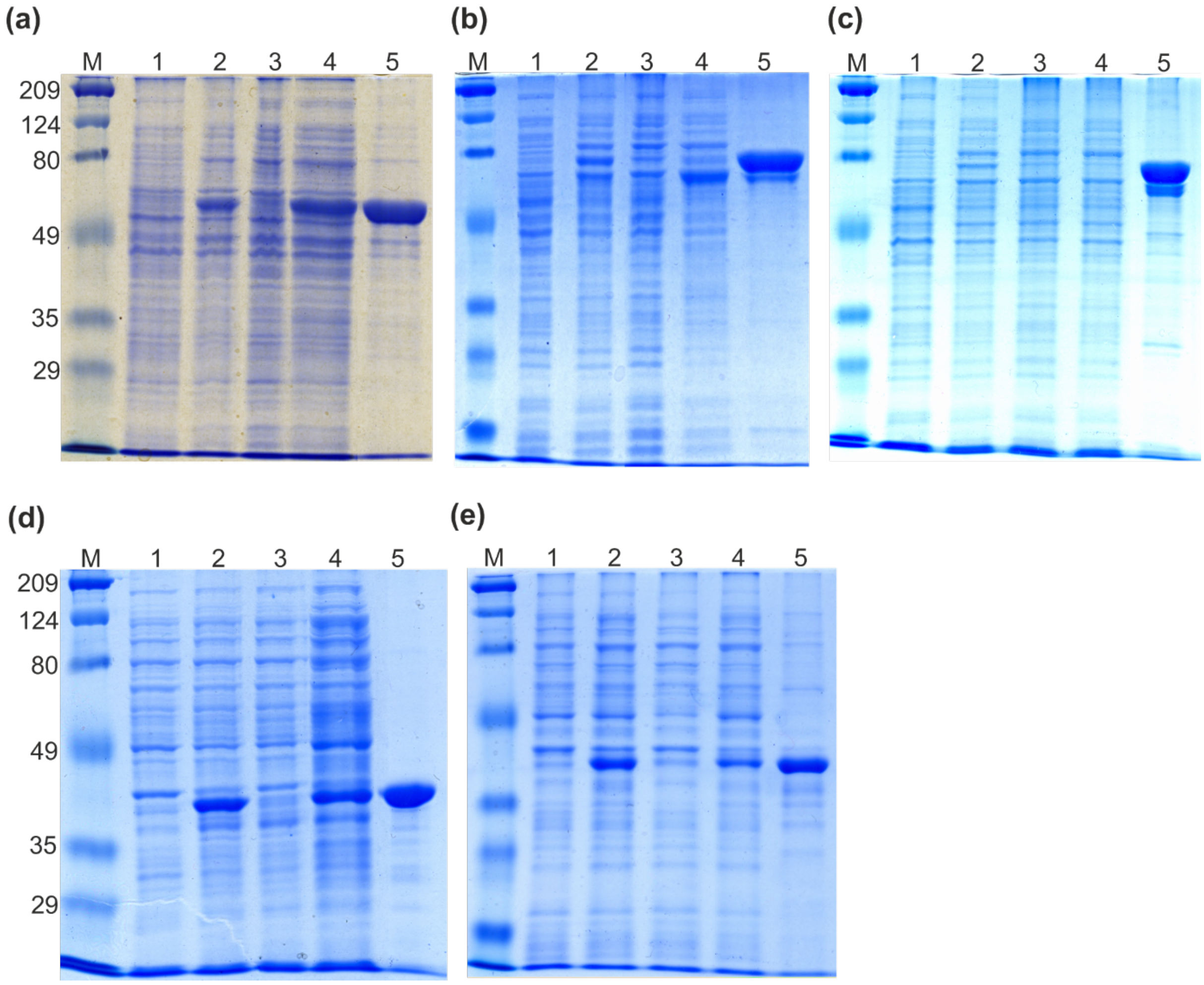


**Figure S3. SDS-polyacrylamide gels with samples from purification of (a) BglC (theoretical Mw = 54.2 kDa), (b) BglC-CtDoc (theoretical Mw = 63.5 kDa), (c) BglC-AcDoc (theoretical Mw = 63.9 kDa), (d) GFP-CtDoc (theoretical Mw = 36.5 kDa), and (e) CFP-AcDoc (theoretical Mw = 37.3 kDa).** Lane 1, cell-free extract prepared from sample collected before induction of gene expression; lane 2, cell-free extract prepared from sample collected at the end of the culture (after induction of gene expression); lane 3, flow-through fraction from NiNTA-based protein purification; lane 4, wash fraction from protein purification; lane 5, sample of purified protein obtained after merging elution fractions with the highest β-glucosidase activity (gels a, b, and c) or elution fractions with the highest fluorescence (gels d and e). Defined quantity of total protein (5 µg) was loaded in each well. M, protein marker (values are in kDa).

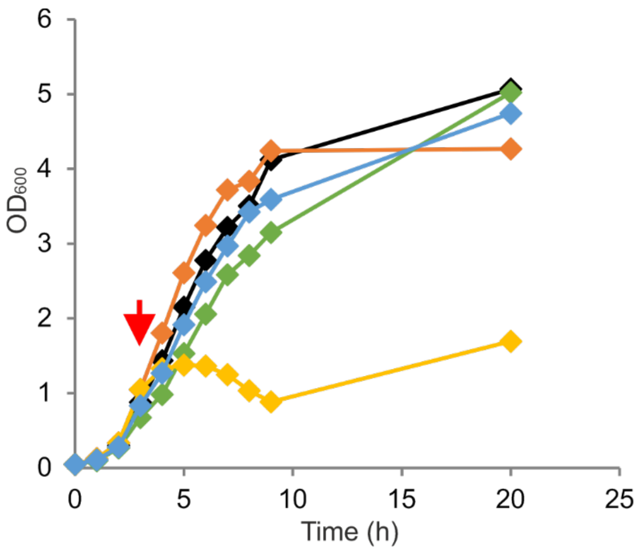


**Figure S4. Effect of an autotransporter expression on viability of the host bacterium *Pseudomonas putida* EM42.** Expression of an autotransporter gene (*estP* in yellow, *int* in green, *igA* in blue and *ag43* in orange) cloned in the pSEVA238 plasmid (control with empty pSEVA238 is shown in black) was induced 3 h after culture start (red arrow) with 1 mM 3-methylbenzoate. Data points show mean values from two biological replicates.

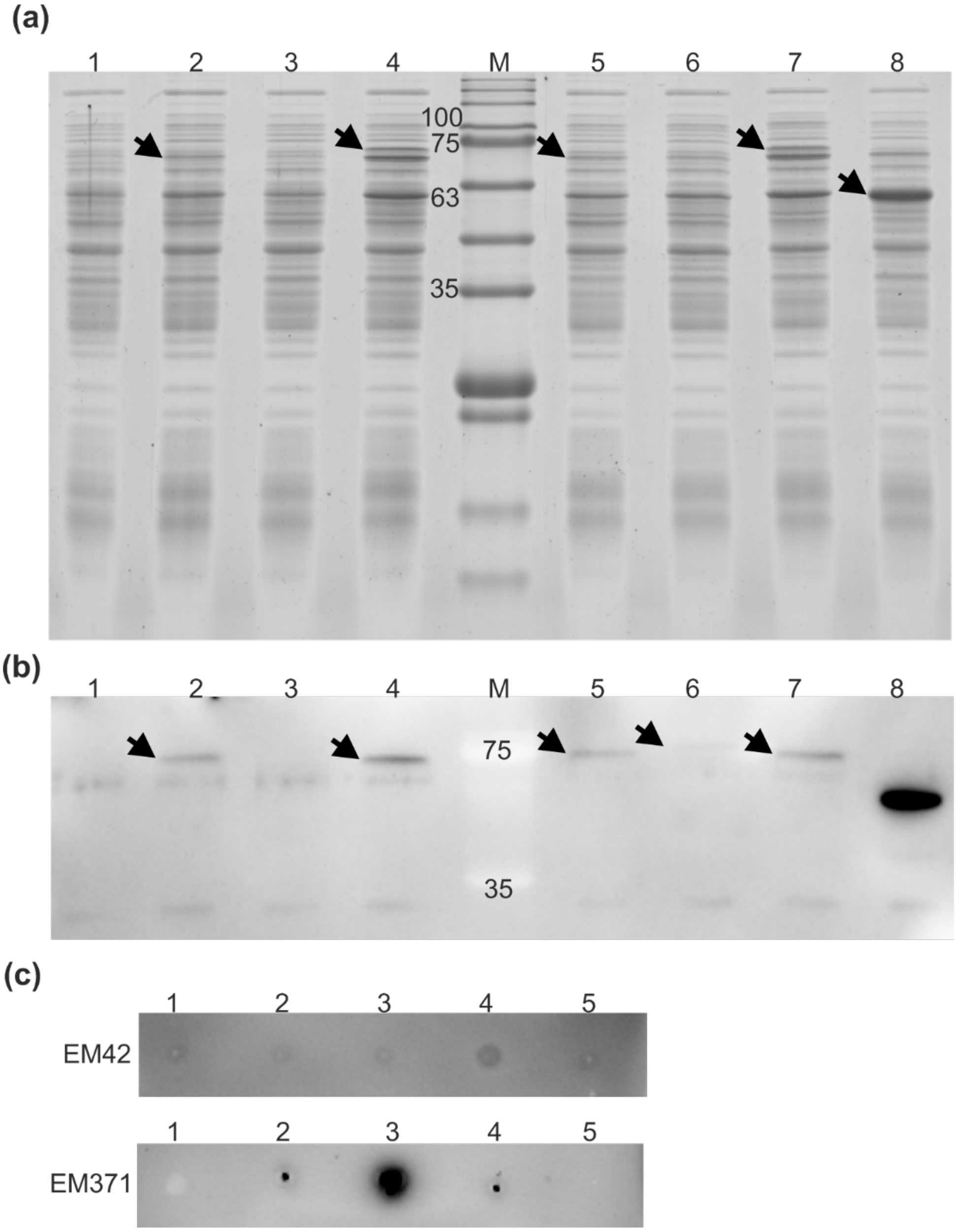


**Figure S5. Expression and display analysis of three different autotransporters with the CtCoh passenger in *Pseudomonas putida* EM42 and EM371.** (a) Sodium dodecyl sulfate polyacrylamide gel electrophoresis (12 % gel) and (b) parallel western blot analysis of cell lysates obtained from *P. putida* EM42 pSEVA238 (1, negative control), *P. putida* EM42 pSEVA238_*igAAT-ctCoh* (2), *P. putida* EM42 pSEVA238_*intAT-ctCoh* (3), *P. putida* EM42 pSEVA238b_*ag43At-ctCoh* (4), *P. putida* EM371 pSEVA238_*igAAT-ctCoh* (5), *P. putida* EM371 pSEVA238_*intAT-ctCoh* (6), *P. putida* EM371 pSEVA238b_*ag43At-ctCoh* (7), and *P. putida* EM42 pSEVA238b_*bglC* (8, positive control) grown in lysogeny broth and induced for 4.5 h with 0.5 mM 3-methylbenzoate. Theoretical molecular weights of detected proteins whose bands are designated with arrows are 67.7 kDa for IgAAT-CtCoh, 87.6 kDa for IntAT-CtCoh, 69.6 kDa for Ag43AT-CtCoh, and 53.4 kDa for BglC control. Equal amounts of total protein (5 μg) were loaded per gel well. M is protein marker, values are in kDa. (c) Dot blot analysis of whole pre-induced *P. putida* EM42 and EM371 cells expressing *igAAT-ctCoh* (1), *intAT-ctCoh* (2), *ag43At-ctCoh* (3), *bglC* (4, cytoplasmic protein control), or bearing empty pSEVA238 plasmid (5, negative control). In case of (b) and (c), proteins with

6xHis tag were detected using mouse anti-His tag monoclonal antibody-HRP conjugate (Thermo Fisher Scientific, USA).

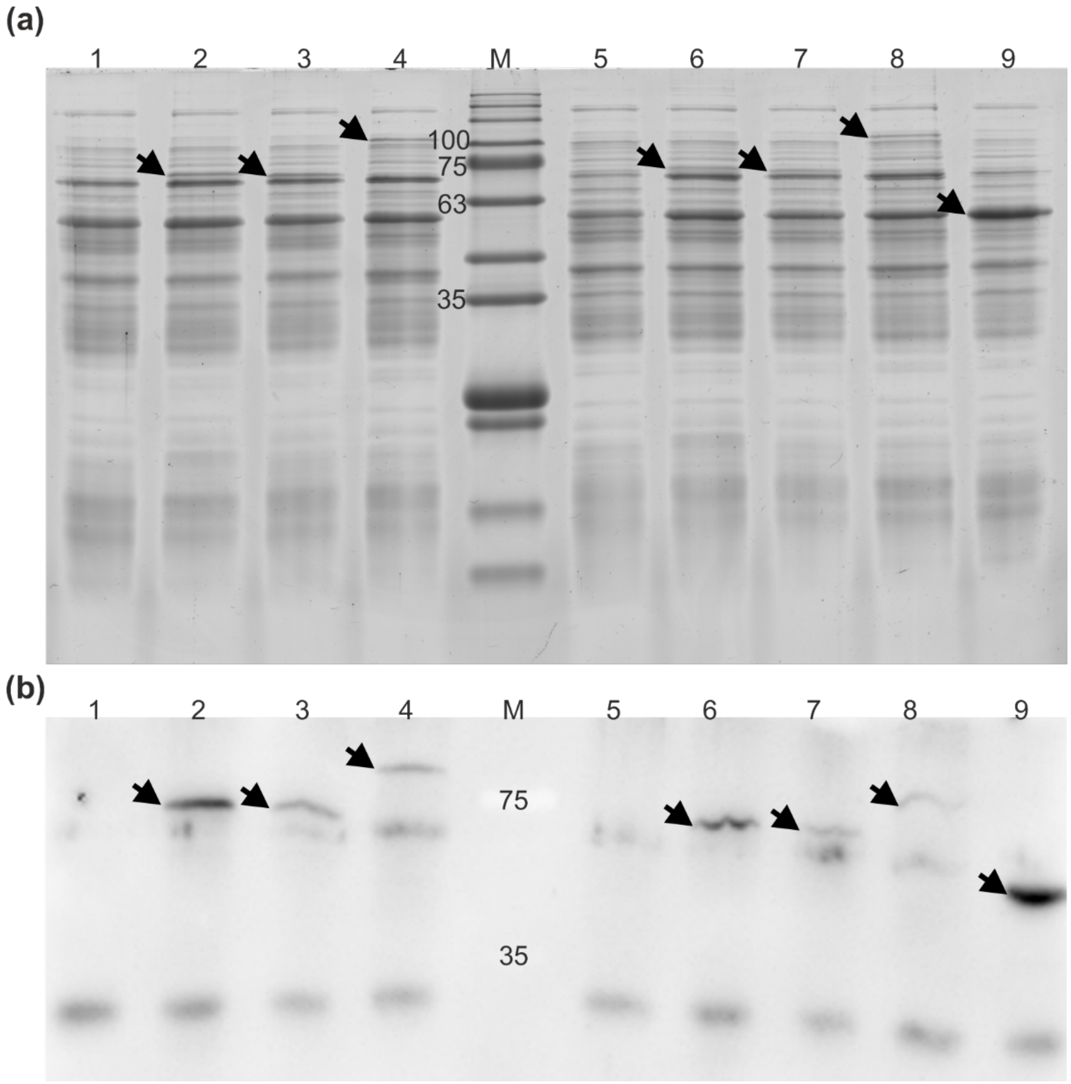


**Figure S6. Expression analysis of three scaffoldin variants displayed *via* Ag43 autotransporter in *Pseudomonas putida* EM42 and EM371.** (a) Sodium dodecyl sulfate polyacrylamide gel electrophoresis (12 % gel) and (b) parallel western blot analysis of cell lysates obtained from *P. putida* EM42 pSEVA238b_*ag43AT* (1, negative control), *P. putida* EM42 pSEVA238b_*ag43AT-acCoh* (2), *P. putida* EM42 pSEVA238b_*ag43AT-ctCoh* (3), *P. putida* EM42 pSEVA238b_*ag43At-acCoh-ctCoh* (4), *P. putida* EM371 pSEVA238b_*ag43AT* (5, negative control), *P. putida* EM371 pSEVA238b_*ag43AT-acCoh* (6), *P. putida* EM371 pSEVA238b_*ag43At-ctCoh* (7), *P. putida* EM371 pSEVA238b_*ag43At-acCoh-ctCoh* (8), and *P. putida* EM42 pSEVA238b_*bglC* (8, positive control) grown in lysogeny broth and induced for 4.5 h with 0.5 mM 3-methylbenzoate. Theoretical molecular weights of detected proteins whose bands are indicated by arrows are 69.9 kDa for Ag43AT-AcCoh, 69.6 kDa for Ag43AT-CtCoh, 88.1 kDa for Ag43AT-AcCoh-CtCoh, and 53.4 kDa for BglC control. Equal amounts of total protein - 5 µg in (a) and 10 µg in (b) - were loaded per gel well. M is protein marker, values are in kDa. In (b), proteins with 6xHis tag were detected using mouse anti-His tag monoclonal antibody-HRP conjugate (Thermo Fisher Scientific, USA).

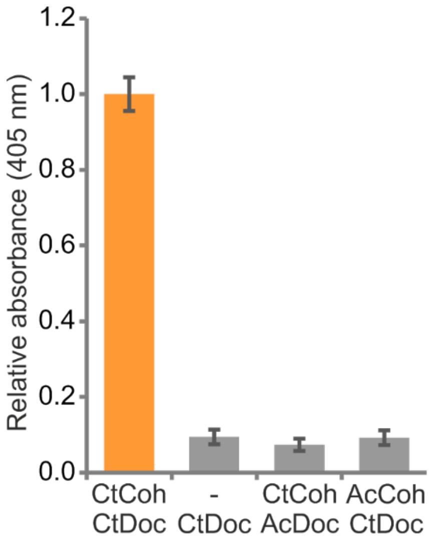


**Figure S7. Cross-reactivity test of the cohesin-dockerin pairs.** *Pseudomonas putida* EM371 cells with the Ag43 autotransporer system displaying CtCoh or AcCoh were mixed with BglC-AcDoc or BglC-CtDoc, respectively, washed and incubated with *p*-nitrophenyl-*β*-D-glucopyranoside. End-point absorbance of the reaction product *p*-nitrophenol (corresponding to β-glucosidase activity of the whole cells with anchored BglC) was determined and compared with the absorbance measured for EM371 pSEVA238b_*Ag43pol* cells mixed with BglC-CtCoh (negative control) and with absorbance of EM371 pSEVA238b_*Ag43AT-ctCoh* cells mixed with BglC-CtCoh (positive control).

### Supplementary sequences

#### *scaf19LKT*

ATGGCAAACACCCCTGTCAGCGGTAACCTCAAGGTCGAGTTCTACAACAGCAACCCGAGCGACACGACGAACAGC
ATCAACCCGCAGTTCAAGGTCACGAACACCGGCAGCAGCGCAATCGACCTCTCCAAGCTGACTCTCCGTTACTAT
TATACCGTCGATGGCCAGAAGGACCAGACCTTCTGGTGCGACCACGCTGCAATCATCGGCAGCAACGGTAGCTAC
AACGGCATCACCTCCAACGTCAAGGGCACCTTCGTCAAGATGTCCTCCTCGACCAACAACGCAGACACCTACCTG
GAGATCAGCAGCACCGGCGGTACGCTGGAGCCTGGTGCACACGTTCAAATCCAGGGTCGTTTCGCAAAGAACGAC
TGGAGCAACTACACCCAGTCGAACGACTACTCGTTCAAGAGCGCCTCGCAGTTCGTCGAGTGGGACCAGGTCACC
GCATACCTGAACGGCGTCCTGGTCTGGGGTAAGGAGCCTGGTGGTAGCGTTGTTCCATCGACTCAGCCAGTTACC
ACCCCACCTGCTACTACCAAGCCTCCTGCTACCACTAAGCCACCAGCTACCACTATCCCGCCATCGGGTTCCGAT
TTGCAGGTGGACATCGGTAGCACTAGCGGTAAGGCTGGTAGCGTAGTGAGCGTTCCAATCACCTTCACCAACGTG
CCGAAGTCGGGCATCTACGCGCTGAGCTTCCGGACTAACTTCGACCCCCAGAAGGTTACCGTGGCTTCCATCGAC
GCCGGCTCGTTGATTGAGAACGCCAGCGATTTCACTACCTACTACAACAACGAGAACGGCTTCGCCTCGATGACC
TTCGAGGCCCCGGTGGACCGGGCCCGCATTATTGACAGCGATGGCGTATTCGCCACTATCAACTTCAAGGTGAGC
GACAGCGCCAAGGTGGGCGAACTGTACAACATCACCACCAACAGCGCCTACACCTCGTTCTACTACTCCGGGACC
GACGAGATCAAGAACGTGGTGTACAACGACGGCAAGATCGAGGTGATCGCCAGCCCGACCCCGACTCAGTCGGCC
ACCCCGACGGTAACCCCGTCGGCCACGGCCACGCCGACGCAATCCGCCACGCCGACCGTAACGCCCTCCGATGGC
GTAGTAGTGGAAATTGGCAAAGTGACGGGCAGCGTGGGCACCACCGTGGAAATTCCGGTGTACTTTCGCGGCGTG
CCCTCCAAAGGCATTGCCAATTGTGATTTTGTGTTTCGCTACGATCCGAATGTGTTGGAAATTATTGGGATTGAT
CCCGGCGATATCATCGTGGACCCGAACCCGACCAAGAGCTTTGATACCGCCATCTACCCGGATCGCAAGATCATC
GTGTTCCTGTTCGCGGAAGACAGCGGCACCGGCGCGTATGCCATTACCAAGGATGGCGTGTTCGCCAAAATCCGC
GCCACCGTGAAATCGTCCGCGCCGGGCTATATTACCTTTGACGAAGTGGGCGGCTTTGCCGATAATGACTTGGTG
GAACAGAAGGTCTCGTTCATCGACGGGGGGGTGAACGTGGGCAACGCCACCCCGACCAAAGGCGCCACCCCCACC
AATACCGCGACGCCGACCAAATCGGCGACGGCGACGCCCACCCGCCCCTCGGCCCCGACGAATACGCCGACCAAT
ACCCCGTCCTCGCCCGGCAATAAAATGAAAATTCAAATCGGGGACGTCAAAGCGAATCAAGGCGACACCGTGATC
GTCCCGATCACCTTTAATGAAGTGCCGGTCATGGGGGTGAATAATTGCAACTTCACCCTGGCGTATGACAAAAAC
ATCATGGAGTTCATCAGCGCGGACGCCGGGGACATCGTCACCCTGCCCATGGCGAACTATAGCTATAACATGCCC
AGCGACGGGCTGGTCAAATTCCTGTATAACGACCAAGCGCAGGGCGCCATGTCGATCAAAGAAGACGGCACCTTC
GCGAACGTGAAGTTCAAGATCAAGCAGTCCGCCGCCTTCGGGAAGTACTCGGTCGGCATCAAGGCCATCGGCTCG
ATCTCCGCGCTGAGCAACTCCAAGCTGATCCCCATCGAATCGATCTTCAAGGACGGCAGCATCACCGTCACCAAC
CTGGAACACCATCACCATCATCACTAA

#### *bglC-ctDoc*

ATGCACCATCACCATCACCATACCTCGCAATCGACGACTCCTCTGGGCAATCTCGAGGAGACTCCCAAACCGGAT
ATCCGCTTCCCGTCCGATTTCGTGTGGGGAGTGGCGACCGCTTCGTTCCAGATCGAAGGCTCCACCACGGCCGAC
GGCCGCGGCCCCAGCATCTGGGACACCTTCTGCGCCACTCCGGGCAAGGTCGAGAACGGCGACACGGGCGACCCT
GCCTGCGACCACTACAACCGGTACCGCGATGACGTGGCCTTGATGCGGGAGCTGGGCGTGGGCGCCTACCGCTTC
TCCATCGCCTGGCCGCGGATCCAGCCCGAGGGCAAGGGCACGCCCGTGGAGGCCGGGCTGGACTTCTACGACCGG
CTTGTGGACTGCCTGCTGGAGGCCGGCATCGAGCCGTGGCCGACCCTCTACCACTGGGACCTGCCGCAGGCGCTG
GAGGACGCGGGCGGCTGGCCCAACCGGGACACGGCCAAGCGGTTCGCCGACTACGCGGAGATCGTCTACCGCCGG
CTCGGCGACCGGATCACCAACTGGAACACGCTCAACGAGCCGTGGTGCTCCGCGTTCCTGGGCTACGCCTCCGGC
GTGCACGCCCCGGGCCGCCAGGAGCCGGCTGCTGCGCTGGCCGCCGCCCACCACCTGATGCTGGGCCACGGGCTG
GCCGCTGCCGTGATGCGGGACTTGGCGGGCCAGGCCGGACGTTCCGTGCGGATCGGTGTCGCGCACAACCAGACC
ACGGTCCGTCCCTACACTGACAGTGAGGCCGACCGGGACGCTGCGCGCCGGATTGACGCCCTGCGGAACCGCATC
TTCACCGAGCCGCTGGTGAAGGGCCGCTACCCGGAGGACCTGATCGAGGACGTCGCCGCGGTCACCGACTACAGC
TTCGTCCAGGACGGCGACCTGAAGACCATCTCCGCCAACCTGGACATGATGGGCGTCAACTTCTACAACCCGAGC
TGGGTGTCAGGCAACCGGGAGAACGGGGGCTCCGACCGGCTGCCCGACGAGGGCTACTCGCCGTCGGTCGGCAGC
GAGCATGTCGTGGAGGTGGACCCCGGCCTGCCGGTGACCGCCATGGGCTGGCCGATCGACCCGACCGGGCTGTAC
GACACGCTGACCCGGCTGGCCAACGACTACCCGGGCCTGCCGCTGTACATCACCGAGAACGGCGCCGCCTTCGAG
GACAAGGTGGTCGACGGCGCGGTGCACGACACCGAGCGGATCGCCTACCTGGACTCGCACCTGCGGGCCGCGCAC
GCTGCCATTGAGGCGGGCGTGCCGCTCAAGGGCTACTTCGCCTGGTCGTTCATGGACAACTTCGAGTGGGCCCTC
GGGTACGGGAAGCGGTTCGGCATCGTGCACGTGGACTACGAGAGCCAGACGCGCACGGTGAAGGACAGCGGCTGG
TGGTACTCCCGGGTGATGCGCAACGGGGGAATCTTCGGACAGGAATCCGGGGGTGGTTCCGCTAGCGGGACGCCC
AGCACCAAACTGTACGGCGATGTCAATGATGACGGCAAAGTGAACTCGACCGACGCCGTCGCCCTGAAGCGGTAT
GTGCTGCGGTCGGGCATCAGCATCAACACCGACAATGCCGACCTGAATGAAGACGGCCGGGTCAATTCGACCGAC
CTGGGCATTCTGAAGCGCTATATTCTCAAAGAAATTGACACGCTGCCGTACAAGAACCACCACCATCACCACCAC
TAA

***bglC-acDoc***
ATGCACCATCACCATCACCATACCTCGCAATCGACGACTCCTCTGGGCAATCTCGAGGAGACTCCCAAACCGGAT
ATCCGCTTCCCGTCCGATTTCGTGTGGGGAGTGGCGACCGCTTCGTTCCAGATCGAAGGCTCCACCACGGCCGAC
GGCCGCGGCCCCAGCATCTGGGACACCTTCTGCGCCACTCCGGGCAAGGTCGAGAACGGCGACACGGGCGACCCT
GCCTGCGACCACTACAACCGGTACCGCGATGACGTGGCCTTGATGCGGGAGCTGGGCGTGGGCGCCTACCGCTTC
TCCATCGCCTGGCCGCGGATCCAGCCCGAGGGCAAGGGCACGCCCGTGGAGGCCGGGCTGGACTTCTACGACCGG
CTTGTGGACTGCCTGCTGGAGGCCGGCATCGAGCCGTGGCCGACCCTCTACCACTGGGACCTGCCGCAGGCGCTG
GAGGACGCGGGCGGCTGGCCCAACCGGGACACGGCCAAGCGGTTCGCCGACTACGCGGAGATCGTCTACCGCCGG
CTCGGCGACCGGATCACCAACTGGAACACGCTCAACGAGCCGTGGTGCTCCGCGTTCCTGGGCTACGCCTCCGGC
GTGCACGCCCCGGGCCGCCAGGAGCCGGCTGCTGCGCTGGCCGCCGCCCACCACCTGATGCTGGGCCACGGGCTG
GCCGCTGCCGTGATGCGGGACTTGGCGGGCCAGGCCGGACGTTCCGTGCGGATCGGTGTCGCGCACAACCAGACC
ACGGTCCGTCCCTACACTGACAGTGAGGCCGACCGGGACGCTGCGCGCCGGATTGACGCCCTGCGGAACCGCATC
TTCACCGAGCCGCTGGTGAAGGGCCGCTACCCGGAGGACCTGATCGAGGACGTCGCCGCGGTCACCGACTACAGC
TTCGTCCAGGACGGCGACCTGAAGACCATCTCCGCCAACCTGGACATGATGGGCGTCAACTTCTACAACCCGAGC
TGGGTGTCAGGCAACCGGGAGAACGGGGGCTCCGACCGGCTGCCCGACGAGGGCTACTCGCCGTCGGTCGGCAGC
GAGCATGTCGTGGAGGTGGACCCCGGCCTGCCGGTGACCGCCATGGGCTGGCCGATCGACCCGACCGGGCTGTAC
GACACGCTGACCCGGCTGGCCAACGACTACCCGGGCCTGCCGCTGTACATCACCGAGAACGGCGCCGCCTTCGAG
GACAAGGTGGTCGACGGCGCGGTGCACGACACCGAGCGGATCGCCTACCTGGACTCGCACCTGCGGGCCGCGCAC
GCTGCCATTGAGGCGGGCGTGCCGCTCAAGGGCTACTTCGCCTGGTCGTTCATGGACAACTTCGAGTGGGCCCTC
GGGTACGGGAAGCGGTTCGGCATCGTGCACGTGGACTACGAGAGCCAGACGCGCACGGTGAAGGACAGCGGCTGG
TGGTACTCCCGGGTGATGCGCAACGGGGGAATCTTCGGACAGGAATCCGGGGGTGGTTCCGCTAGCAAGTTCATC
TACGGCGACGTGGACGGCAACGGCAGCGTCCGCATCAACGACGCTGTGCTGATCCGTGACTACGTTCTGGGCAAG
ATCAACGAGTTCCCCTACGAATATGGTATGTTAGCTGCTGATGTAGATGGTAACGGCTCCATCAAGATCAACGAC
GCCGTCCTGGTGCGCGACTACGTGCTGGGCAAGATCTTCCTGTTCCCCGTCGAAGAAAAGGAAGAACACCACCAT
CACCATCACTAA

***gfp-ctDoc***
ATGAGTAAAGGAGAAGAACTTTTCACCGGTGTTGTTCCGATCCTGGTTGAACTGGATGGTGATGTTAACGGCCAC
AAATTCTCTGTTCGTGGTGAAGGTGAAGGTGATGCAACCAACGGTAAACTGACCCTGAAATTCATCTGCACTACC
GGTAAACTGCCGGTTCCATGGCCGACTCTGGTGACTACCCTGACCTATGGTGTTCAGTGTTTTTCTCGTTACCCG
GATCACATGAAGCAGCATGATTTCTTCAAATCTGCAATGCCGGAAGGTTATGTACAGGAGCGCACCATTTCTTTC
AAAGACGATGGCACCTACAAAACCCGTGCAGAGGTTAAATTTGAAGGTGATACTCTGGTGAACCGTATTGAACTG
AAAGGCATTGATTTCAAAGAGGACGGCAACATCCTGGGCCACAAACTGGAATATAACTTCAACTCCCATAACGTT
TACATCACCGCAGACAAACAGAAGAACGGTATCAAAGCTAACTTCAAAATTCGCCATAACGTTGAAGACGGTAGC
GTACAGCTGGCGGACCACTACCAGCAGAACACTCCGATCGGTGATGGTCCGGTTCTGCTGCCGGATAACCACTAC
CTGTCCACCCAGTCTAAACTGTCCAAAGACCCGAACGAAAAGCGCGACCACATGGTGCTGCTGGAGTTCGTTACT
GCCGCAGGTATCACGCACGGCATGGATGAACTATACAAAGAATTCGAGCTCTCCGGGGGTGGTTCCGCTAGCGGG
ACGCCCAGCACCAAACTGTACGGCGATGTCAATGATGACGGCAAAGTGAACTCGACCGACGCCGTCGCCCTGAAG
CGGTATGTGCTGCGGTCGGGCATCAGCATCAACACCGACAATGCCGACCTGAATGAAGACGGCCGGGTCAATTCG
ACCGACCTGGGCATTCTGAAGCGCTATATTCTCAAAGAAATTGACACGCTGCCGTACAAGAACCACCACCATCAC
CACCACTAA

***cfp-acDoc***
ATGGGCCATCATCATCACCATCACGTGAGCAAGGGCGAGGAGCTGTTCACCGGGGTGGTGCCCATCCTGGTCGAG
CTGGACGGCGACGTAAACGGCCACAAGTTCAGCGTGTCCGGCGAGGGCGAGGGCGATGCCACCTACGGCAAGCTG
ACCCTGAAGTTCATCTGCACCACCGGCAAGCTGCCCGTGCCCTGGCCCACCCTCGTGACCACCCTGACCTGGGGC
GTGCAGTGCTTCGCCCGCTACCCCGACCACATGAAGCAGCACGACTTCTTCAAGTCCGCCATGCCCGAAGGCTAC
GTCCAGGAGCGCACCATCTTCTTCAAGGACGACGGCAACTACAAGACCCGCGCCGAGGTGAAGTTCGAGGGCGAC
ACCCTGGTGAACCGCATCGAGCTGAAGGGCATCGACTTCAAGGAGGACGGCAACATCCTGGGGCACAAGCTGGAG
TACAACGCCATCAGCGACAACGTCTATATCACCGCCGACAAGCAGAAGAACGGCATCAAGGCCAACTTCAAGATC
CGCCACAACATCGAGGACGGCAGCGTGCAGCTCGCCGACCACTACCAGCAGAACACCCCCATCGGCGACGGCCCC
GTGCTGCTGCCCGACAACCACTACCTGAGCACCCAGTCCAAGCTGAGCAAAGACCCCAACGAGAAGCGCGATCAC
ATGGTCCTGCTGGAGTTCGTGACCGCCGCCGGGATCACTCTCGGCATGGACGAGCTGTACAAGGGTACCAAATTT
ATATATGGTGATGTTGATGGTAATGGAAGTGTAAGAATTAATGATGCTGTCCTAATAAGAGACTATGTATTAGGA
AAAATCAATGAATTCCCATATGAATATGGTATGCTTGCAGCAGATGTTGATGGTAATGGAAGTATAAAAATTAAT
GATGCTGTTCTAGTAAGAGACTACGTGTTAGGAAAGATATTTTTATTCCCTGTTGAAGAGAAAGAAGAACTCGAG
CACCACCACCACCACCACTGA

***igAAT***

ATGAAATACCTATTGCCTACGGCAGCCGCTGGATTGTTATTACTCGCCGCCCAGCCGGCCATGGCGATCGTTCTT
GGTGCGCCGGTGCCGTATCCCGATCCGCTGGAACCGCGTGCCGCCGCCGCGGCCGCGAATTCGAGCTCGGTACCC
GGGGATCCTCTAGAGTCGACCTGCAGGCATGCAAGCTTGCGGCCGCTCGACAATTCAGCCGCAATTAGTATGGCA
AATCCACGTCCACCAACACCGCGGGCTGCTGCGGCCGTATTTTCATTGGATGATTATGATGCAAAAGACAATAGT
GAATCATCAATAGGTAATTTAGCTCGTGTAATACCTAGAATGGGAAGGGAGCTAATTAATGATTATGAAGAAATC
CCCTTGGAGGAGTTGGAAGATGAAGCGGAAGAAGAACGTCGCCAAGCAACGCAATTCCAACCCAAAAGTCGTAAC
CGTAGAGCTATATCATCGGAACCATCATCTGATGAAGATGCATCTGAATCGGTTTCCACATCAGACAAACACCCT
CAAGATAATACGGAACTTCATGAAAAAGTTGAGACGGCGGGTTTACAACCAAGAGCCGCGCAGCCGCGAACCCAA
GCCGCCGCGCAAGCCGATGCAGTCAGCACCAATACTAACTCGGCTTTATCTGACGCAATGGCAAGCACGCAATCT
ATCTTGTTGGATACAGGTGCTTCATTAACACGGCACATTGCACAAAAATCACGCGCTGATGCCGAAAAAAACAGT
GTTTGGATGTCAAACACCGGTTATGGCCGTGATTATGCTTCCGCACAATATCGCCGGTTTAGTTCGAAACGCACG
CAAACACAAATCGGCATTGACCGCAGCTTGTCCGAAAATATGCAGATAGGCGGAGTATTGACTTACTCTGACAGT
CAGCATTCTTTTGATCTGGCGGGCGGCAAAAATACTTTTGTGCAAGCCAACCTTTATGGTAAGTATTATCTAAAT
GATGCTTGGTATGTGGCCGGCGATATTGGTGCGGGCAGCTTGAGAAGCCGGTTACAAACGCAGCAAAAAGCAAAC
TTTAACCGAACAAGCATCCAAACCGGCCTTACTTTGGGCAATACGCTGAAAATCAATCAATTCGAGATTGTCCCT
AGTGCGGGTATCCGTTACAGCCGCCTGTCATCTGCTGATTACAAGTTGGGTGACGACAGTGTTAAAGTAAGTTCT
ATGGCAGTGAAAACACTAACGGCCGGACTGGATTTTGCTTATCGGTTTAAAGTCGGCAACCTTACCGTAAAACCC
TTGTTATCTGCTGCTTACTTTGCCAATTATGGCAAAGGCGGCGTGAATGTGGGCGGTAAATCCTTCGCCTATAAA
GCAGATAATCAACAGCAATATTCAGCAGGCGCCGCGTTACTGTACCGTAATGTTACATTAAACGTAAATGGCAGT
ATTACAAAAGGAAAACAATTGGAAAAACAAAAATCCGGACAAATTAAAATACAGATTCGTTTCTAA

***intAT***

ATGATTACTCATGGTTGTTATACCCGGACCCGGCACAAGCATAAGCTAAAAAAAACATTGATTATGCTTAGTGCT
GGTTTAGGATTGTTTTTTTATGTTAATCAGAACTCATTTGCAAATGGTGAAAATTATTTTAAATTGGGTTCGGAT
TCAAAACTGTTAACTCATGATAGCTATCAGAATCGCCTTTTTTATACGTTGAAAACTGGTGAAACTGTTGCCGAT
CTTTCTAAATCGCAAGATATTAATTTATCGACGATTTGGTCGTTGAATAAGCATTTATACAGTTCTGAAAGCGAA
ATGATGAAGGCCGCGCCTGGTCAGCAGATCATTTTGCCACTCAAAAAACTTCCCTTTGAATACAGTGCACTACCA
CTTTTAGGTTCGGCACCTCTTGTTGCTGCTGGTGGTGTTGCTGGTCACACGAATAAACTGACTAAAATGTCCCCG
GACGTGACCAAAAGCAACATGACCGATGACAAGGCATTAAATTATGCGGCACAACAGGCGGCGAGTCTCGGTAGC
CAGCTTCAGTCGCGATCTCTGAACGGCGATTACGCGAAAGATACCGCTCTTGGTATCGCTGGTAACCAGGCTTCG
TCACAGTTGCAGGCCTGGTTACAACATTATGGAACGGCAGAGGTTAATCTCCAGAGTGGTAATAACTTTGACGGT
AGTTCACTGGACTTCTTATTACCGTTCTATGATTCCGAAAAAATGCTGGCATTTGGTCAGGTCGGAGCGCGTTAC
ATTGACTCCCGCTTTACGGCAAATTTAGGTGCGGGTCAGCGTTTTTTCCTTCCTGCAAACATGTTGGGCTATAAC
GTCTTCATTGATCAGGATTTTTCTGGTGATAATACCCGTTTAGGTATTGGTGGCGAATACTGGCGAGACTATTTC
AAAAGTAGCGTTAACGGCTATTTCCGCATGAGCGGCTGGCATGAGTCATACAATAAGAAAGACTATGATGAGCGC
CCAGCAAATGGCTTCGATATCCGTTTTAATGGCTATCTACCGTCATATCCGGCATTAGGCGCCAAGCTGATATAT
GAGCAGTATTATGGTGATAATGTTGCTTTGTTTAATTCTGATAAGCTCCAGTCGAATCCTGGTGCGGCGACCGTT
GGTGTAAACTATACTCCGATTCCTCTGGTGACGATGGGGATCGATTACCGTCATGGTACGGGTAATGAAAATGAT
CTCCTTTACTCAATGCAGTTCCGTTATCAGTTTGATAAATCGTGGTCTCAGCAAATTGAACCACAGTATGTTAAC
GAGTTAAGAACATTATCAGGCAGCCGTTACGATCTGGTTCAGCGTAATAACAATATTATTCTGGAGTACAAGAAG
CAGGATATTCTTTCTCTGAATATTCCGCATGATATTAATGGTACTGAACACAGTACGCAGAAGATTCAGTTGATC
GTTAAGAGCAAATACGGTCTGGATCGTATCGTCTGGGATGATAGTGCATTACGCAGTCAGGGCGGTCAGATTCAG
CATAGCGGAAGCCAAAGCGCACAAGACTACCAGGCTATTTTGCCTGCTTATGTGCAAGGTGGCAGCAATATTTAT
AAAGTGACGGCTCGCGCCTATGACCGTAATGGCAATAGCTCTAACAATGTACAGCTTACTATTACCGTTCTGTCG
AATGGTCAAGTTGTTGACCAGGTTGGGGTAACGGACTTTACGGCGGATAAGACTTCGGCTAAAGCGGATAACGCC
GATACCATTACTTATACCGCGACGGTGAAAAAGAATGGGGTAGCTCAGGCTAATGTCCCTGTTTCATTTAATATT
GTTTCAGGAACTGCAACTCTTGGGGCAAATAGTGCCAAAACGGATGCTAACGGTAAGGCAACCGTAACGTTGAAG
TCGAGTACGCCAGGACAGGTCGTCGTGTCTGCTAAAACCGCGGAGATGACTTCAGCACTTAATGCCAGTGCGGTT
ATATTTTTTGATGCGGCCGCGAATTCGAGCTCGGTACCCGGGGATCCTCTAGAGTCGACCTGCAGGCATGCAAGC
TTGCGGCCGCCCGGTGCGCCGGTGCCGTATCCCGATCCGCTGGAACCGCGTTAA

***ag43AT***
ATGAAACGACATCTGAATACCTGCTACAGGCTGGTATGGAATCACATGACGGGCGCTTTCGTGGTTGCCTCCGAA
CTGGCCCGCGCACGGGGTAAACGTGGCGGTGTGGCGGTTGCACCGTCTCTTGCCGCAGTCACGTCACTCCCGGTG
CTGGCTGCTGACAGATCTGCCATGGCGGAATTCGAGCTCGGTACCCTCGAGCTGAACGGATCCATTGACCCCACG
AATGTCACTCTCGCCTCCGGTGCCACCTGGAATATCCCCGATAACGCCACGGTGCAGTCGGTGGTGGATGACCTC
AGCCATGCCGGACAGATTCATTTCACCTCCACCCGCACAGGGAAGTTCGTACCGGCAACCCTGAAAGTGAAAAAC
CTGAACGGACAGAATGGCACCATCAGCCTGCGTGTACGCCCGGATATGGCACAGAACAATGCTGACAGACTGGTC
ATTGACGGCGGCAGGGCAACCGGAAAAACCATCCTGAACCTGGTGAACGCCGGCAACAGTGCGTCGGGGCTGGCG
ACCAGCGGTAAGGGTATTCAGGTGGTGGAAGCCATTAACGGTGCCACCACGGAGGAAGGGGCCTTTGTCCAGGGG
AACAGGCTGCAGGCCGGTGCCTTTAACTACTCCCTCAACCGGGACAGTGATGAGAGCTGGTATCTGCGCAGTGAA
AATGCTTATCGTGCAGAAGTCCCCCTGTATGCCTCCGTGCTGACACAGGCAATGGACTATGACCGGATTGTGGCA
GGCTCCCGCAGCCATCAGACCGGTGTAAATGGTGAAAACAACAGCGTCCGTCTCAGCATTCAGGGCGGTCATCTC
GGTCACGATAACAATGGCGGTATTGCCCGTGGGGCCACGCCGGAAAGCAGCGGCAGCTATGGATTCGTCCGTCTG
GAGGGTGACCTGATGAGAACAGAGGTTGCCGGTATGTCTGTGACCGCGGGGGTATATGGTGCTGCTGGCCATTCT
TCCGTTGATGTTAAGGATGATGACGGCTCCCGTGCCGGCACGGTCCGGGATGATGCCGGCAGCCTGGGCGGATAC
CTGAATCTGGTACACACGTCCTCCGGCCTGTGGGCTGACATTGTGGCACAGGGAACCCGCCACAGCATGAAAGCG
TCATCGGACAATAACGACTTCCGCGCCCGGGGCTGGGGCTGGCTGGGCTCACTGGAAACCGGTCTGCCCTTCAGT
ATCACTGACAACCTGATGCTGGAGCCACAACTGCAGTATACCTGGCAGGGACTTTCCCTGGATGACGGTAAGGAC
AACGCCGGTTATGTGAAGTTCGGGCATGGCAGTGCACAGCATGTGCGTGCCGGTTTCCGTCTGGGCAGCCACAAC
GATATGACCTTTGGCGAAGGCACCTCATCCCGTGCCCCCCTGCGTGACAGTGCAAAACACAGTGTGAGTGAATTA
CCGGTGAACTGGTGGGTACAGCCTTCTGTTATCCGCACCTTCAGCTCCCGGGGAGATATGCGTGTGGGGACTTCC
ACTGCAGGCAGCGGGATGACGTTCTCTCCCTCACAGAATGGCACATCACTGGACCTGCAGGCCGGACTGGAAGCC
CGTGTCCGGGAAAATATCACCCTGGGCGTTCAGGCCGGTTATGCCCACAGCGTCAGCGGCAGCAGCGCTGAAGGG
TATAACGGTCAGGCCACACTGAATGTGACCTTCTGA

***estPAT***
ATGCGAAAAGCCCCGTTATTGCGCTTTACCCTCGCTTCACTGGCCCTGGCCTGTAGCCAGGCGTTGGCCGGTGCG
CCGGTGCCGTATCCGGACCCGCTGGAACCGCGTGGATCCGTCGACTCTAGACTGCAGTCGGTCCACCCGACCATC
GCGGGTCAGCAGCTGATTGCCGATTACGCCTACTCGATCCTCGCGGCCCCCTGGGAACTGACCCTGCTACCGGAA
ATGGCCCACGCCAGCCTGCGGGCTCACCAGGATGAGTTGCGTAATCAGTGGCAGACGCCTTGGCAAGCAGTTGGC
CAATGGCAAGCCTTTGTCGCCAGCGGCGCTCAGGACCTGGACTTCGACGGCCAGCACAGCGCGGCCAGCGGTGAC
GGCCGCGGCTACAACCTGACCGTGGGCGGCAGCTATCGCCTGAACGACGCCTGGCGCCTGGGCCTGGCCGGCGGT
GCAAACCGGCAGAAGCTGGAAGCTGGTGAACAGGACTCGGACTACAAGCTGAACAGTTATATGGCCAGTGCCTTT
GCCCAATACCGCCAGGACCGCTGGTGGGCCGACGCGGCGCTGACCGCCGGGCACCTGGATTACAGCGACCTCAAG
CGTACCTTCGCCCTGGGCGTGAATGACCGCAGTGAGAAGGGCGACACCGACGGCGAGGCCTGGGCAATGTCCGGG
CGGCTGGGCTACAACCTGGCGGCCGACACCAGCAACTGGCAGTTGGCACCTTTCATCAGTGCCGACTATGCGCGG
GTGAAGGTGGATGGCTACGACGAGAAGAGCGGACGTTCGACGGCGCTTGGCTTCGATGACCAGGAGCGCACGTCA
CGCCGCCTGGGCGTGGGGCTGCTGGGCAGTGTGCAGGTACTGCCAAGTACCCGGCTTTTCGCCGAGGTGGCGCAG
GAGCATGAGTTCGAGGACGACGAGCAGGATGTGACGATGCACCTGACCAGCTTGCCGGCGAATGACTTCACCCTG
ACCGGGTATACGCCGCACAGCGACCTGACCCGGGCGAGCCTGGGTGTGAGCCATGAACTGGTGGCAGGGGTGCAT
TTGCGCGGGAACTACAACTGGCGCAAGAGTGATGAGTTGACGCAACAGGGTATTAGCGTGGGGGTTAGCGTGGAC
TTCTGA

### References


(1) Grant, S. G.; Jessee, J.; Bloom, F. R.; Hanahan, D. Differential Plasmid Rescue from Transgenic Mouse DNAs into *Escherichia coli* Methylation-Restriction Mutants. *Proc. Natl. Acad. Sci. U. S. A.* **1990**, *87* (12), 4645–4649.

(2) Manoil, C.; Beckwith, J. TnphoA: A Transposon Probe for Protein Export Signals. *Proc. Natl. Acad. Sci. U. S. A.* **1985**, *82* (23), 8129–8133.

(3) Boyer, H. W.; Roulland-Dussoix, D. A Complementation Analysis of the Restriction and Modification of DNA in *Escherichia coli. J. Mol. Biol.* **1969**, *41* (3), 459–472.

(4) Martínez-García, E.; Nikel, P. I.; Aparicio, T.; de Lorenzo, V. Pseudomonas 2.0: Genetic Upgrading of *P. putida* KT2440 as an Enhanced Host for Heterologous Gene Expression. *Microb. Cell Factories* **2014**, *13*, 159.

(5) Martínez-García, E.; Fraile, S.; Rodríguez Espeso, D.; Vecchietti, D.; Bertoni, G.; de Lorenzo, V. The Naked Cell: Emerging Properties of a Surfome-Streamlined *Pseudomonas putida* Strain. 2020, In preparation.

(6) Kessler, B.; de Lorenzo, V.; Timmis, K. N. A General System to Integrate LacZ Fusions into the Chromosomes of Gram-Negative Eubacteria: Regulation of the Pm Promoter of the TOL Plasmid Studied with All Controlling Elements in Monocopy. *Mol. Gen. Genet. MGG* **1992**, *233* (1–2), 293–301.

(7) Silva-Rocha, R.; Martínez-García, E.; Calles, B.; Chavarría, M.; Arce-Rodríguez, A.; de Las Heras, A.; Páez-Espino, A. D.; Durante-Rodríguez, G.; Kim, J.; Nikel, P. I.; Platero, R.; de Lorenzo, V. The Standard European Vector Architecture (SEVA): A Coherent Platform for the Analysis and Deployment of Complex Prokaryotic Phenotypes. *Nucleic Acids Res.* **2013**, *41* (Database issue), D666-675.

(8) Dvořák, P.; de Lorenzo, V. Refactoring the Upper Sugar Metabolism of *Pseudomonas putida* for Co-Utilization of Cellobiose, Xylose, and Glucose. *Metab. Eng.* **2018**, *48*, 94–108.

(9) Vazana, Y.; Barak, Y.; Unger, T.; Peleg, Y.; Shamshoum, M.; Ben-Yehezkel, T.; Mazor, Y.; Shapiro, E.; Lamed, R.; Bayer, E. A. A Synthetic Biology Approach for Evaluating the Functional Contribution of Designer Cellulosome Components to Deconstruction of Cellulosic Substrates. *Biotechnol. Biofuels* **2013**, *6* (1), 182.

(10) Muñoz-Gutiérrez, I.; Moss-Acosta, C.; Trujillo-Martinez, B.; Gosset, G.; Martinez, A. Ag43-Mediated Display of a Thermostable β-Glucosidase in *Escherichia coli* and Its Use for Simultaneous Saccharification and Fermentation at High Temperatures. *Microb. Cell Factories* **2014**, *13*, 106.
